# Core enhancers of the 3’RR optimize *IgH* nuclear position and loop conformation for oriented CSR

**DOI:** 10.1101/2023.07.11.548507

**Authors:** Charlotte Bruzeau, Justine Pollet, Morgane Thomas, Zhaoqing Ba, David Roulois, Eric Pinaud, Sandrine Le Noir

## Abstract

Class switch recombination is an essential process which enabling B cells to adapt immunoglobulin subtypes to antigens. Transcription plays a crucial role in regulating CSR in which the *IgH 3’Regulatory Region* (*3’RR*) was identified as a key player. The *3’RR* stands at the 3’ end of *IgH* locus and is composed of four core enhancers surrounded by inverted repeated sequences, forming a quasi-palindrome. In addition to transcriptional control, nuclear organization appears to be an important level in CSR regulation. Furthermore, the chromatin loops at *IgH* locus facilitate an efficient CSR recombination by bringing the donor and acceptor switch regions closer together. However, the precise control mechanisms governing both of these processes remain partially understood. Here, using the reference DNA 3D-FISH technique combined with various high throughput approaches, we showed that 3’RR core enhancers are necessary and sufficient to preorganize resting B cell nuclei to facilitate a deletional CSR mechanism at activated stage. We demonstrated that the 3’RR core enhancers regulate *IgH* locus addressing in the nuclei, control *IgH* locus accessibility and orchestrate *IgH* loops formation. Our findings pinpointed an additional regulation level of mechanisms underlying B cell diversification.

**Graphical abstract:** 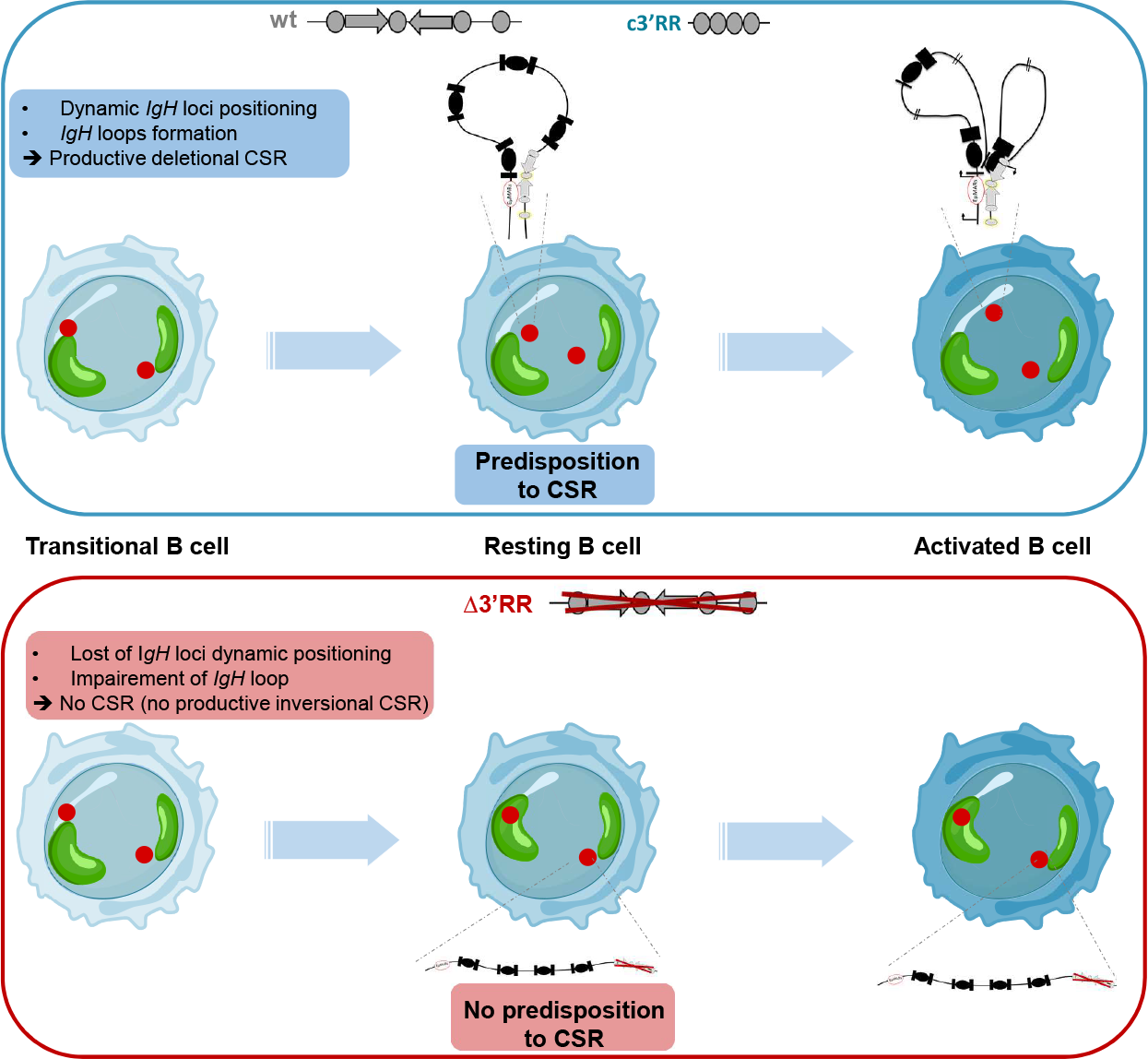

Position of *IgH* loci through B cell development (from transitional to stimulated stages) is represented by the red spots. In *wt* and *c3’RR* context, *IgH* loci get closer from each other and from nucleus center during evolution from transitional to mature resting stage and they relocates more at nuclear periphery, away one from each other, upon *in vitro* stimulation. In *Δ3’RR* model, this dynamic is lost and, moreover, *IgH* loci are more localized to pericentromeric heterochromatin (represented by green area) since the mature resting B cell stage and remain in after *in vitro* stimulation.

## Introduction

Late stages of B cell development are characterized by immunoglobulin genes remodeling to produce highly specific antibodies. Somatic hypermutation (SHM) and class switch recombination (CSR) are the two mains Immunoglobulin Heavy Chain (*IgH*) locus modifications that both require transcription of target sequences and cytidine deamination by the activation-induced cytidine deaminase (AID) enzyme (1). It has been extensively documented in literature that these events are under the transcriptional control of the super-enhancer located at *IgH* locus 3’end: the *IgH 3’Regulatory Region* (*3’RR*) (2). In mice, this region is composed of four core enhancers, (*hs3a, hs1*.*2, hs3b* and *hs4*), which are surrounded by inverted repeated sequence (IRIS) centered on *hs1*.*2* enhancer and delimited by *hs3a* and *hs3b* (3) (Figure 1A). *IgH* locus forms a topologically associated domain (TAD) as demonstrated by chromosome conformation capture (3C)-based studies (4). The 3’ end of the locus is characterized by an insulator region composed of 10 CTCF (CCCTC-binding factor) binding site named 3’ CTCF binding elements (3’CBE) which delimited the 3’ *IgH* TAD border (5). Several mouse models carrying partial or total deletions of *3’RR* region allow to decipher the exact role of both core enhancers and quasi-palindromic structure on SHM, CSR, transcription and immunoglobulin production during B cell development (3). The *3’RR* entire deletion leads to a 2-fold decrease of *IgH* germline transcription (GLT), and consequently, totally abolishes SHM and CSR events (6). Interestingly *3’RR* core enhancers (*c3’RR* mouse model) are mandatory for full activity of the *3’RR* super-enhancer, and the surrounding sequences do not constitute useless junk DNA, but contribute to *3’RR* activity: while CSR is supported by enhancers alone, SHM requires both *3’RR* enhancers and its palindromic architecture (7, 8).

**Figure 1.**
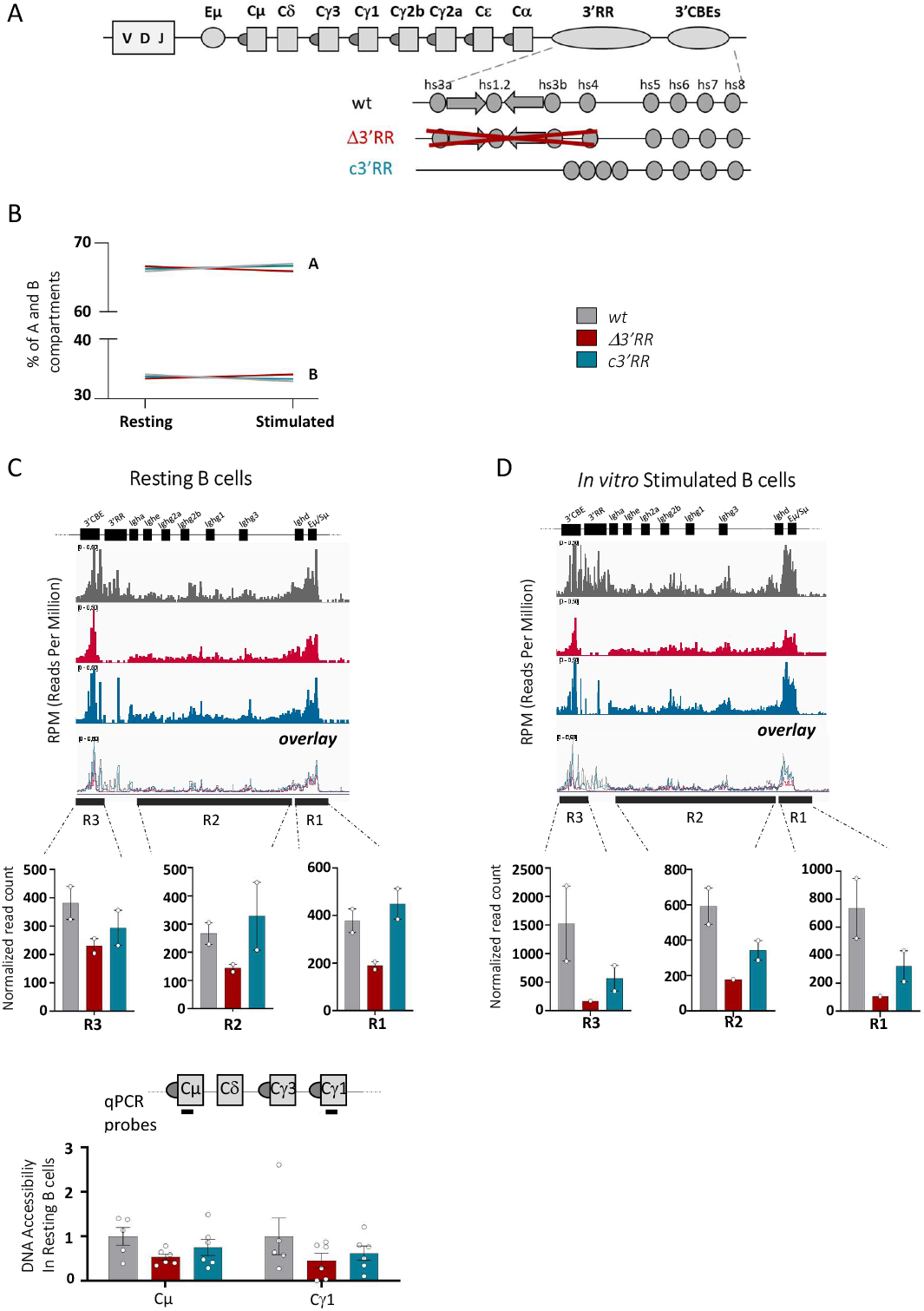
3’RR core enhancers maintained *IgH* loci in an active compartment in mature resting B cells. **A**. Murine *IgH* locus scheme (not to scale). IgH constant genes are represented by squares, arrows represent the palindromic structure. Deletion of *Δ3’RR* (6, 30) *and c’3RR* (7) mice used in this study are schemed. **B**. A and B compartment proportions in resting and stimulated B cells from *wt* and mutants **C**. *IgH* locus accessibility by ATAC-Seq in splenic resting B cells from *wt* (n=2 independent mice), *3’RR* (n=2 independent mice) and *c’3RR* mice (n=2 independent mice). Bedgraphs are normalized to reads per million and visualized with IGV. *IgH* loci was split into three regions and the normalized reads coverage were counted in each region as indicated in each bargraph. Relative DNA accessibility by qPCR in resting B cells. Normalization was done to *Lig1* which is not differentially opened between models. q-PCR probes are indicated with black lines. **D**. As in B for *in vitro* LPS activated B cell from *wt* (n=2 independent mice), *Δ3’RR* (n=1 mouse) and *c3’RR* (n=2 independent mice).

Beyond transcriptional control of SHM and CSR, additional levels of regulation occurred during late B cell maturation. Most of Ig gene regulation studies have been so far performed at nucleosomal scale (epigenetic modifications and regulatory transcription of loci and gene segments). The increasing interest for the understanding of gene regulation at the whole nuclei scale raises the necessity to revisit previous models at both supranucleosomal (DNA loops and A/B compartments) and nuclear (chromosome territories and gene positioning) levels (9). To date, numerous studies showed that nuclear positioning of *IgH* loci is dynamic and regulated during early and late B cell development and *IgH* remodeling regulation, especially CSR recombination, also occurs at supranucleosomal level (10). CSR mechanism involves DNA double strand breaks (DSB) at *Sμ* donor and *Sx* acceptor regions (11). Mammalian switch (*S)* regions, located upstream of each constant gene except Cδ, consist of 1 to 10 kb-long highly repetitive G-rich DNA sequences containing clusters of RGYW AID deamination hotspots (12). AID-induced DSB at *S* regions are followed by the deletion of DNA segment located between both DNA DSB and finally replacing *Cμ* constant gene by one of the six *Cx* constant gene *(IgG3, IgG1, IgG2a, IgG2b, IgA, IgE)*. The recombination event between DNA ends is essentially ensured by the NHEJ complex to achieve productive deletional CSR (13). Mature resting B cells ensure an *IgH* locus configuration “prepared for CSR” by bringing in close contact the *3’RR* and *Eμ*-regions, thereby forming a 50-200Kb “big chromatin loop” that contains all S regions. Upon stimulation, this configuration acquires additional contact between the acceptor S region involved in CSR and the previously interacting *IgH* enhancers (14–16). In this new layout, S donor and S acceptor regions are aligned to favor productive deletional CSR. DSB response factors inhibition impact the ratio of deletional *vs* inversional CSR pathway, highlighting the important role of DNA repair pathway for efficient deletional CSR (17). The current model proposes that such loops are formed by loop extrusion, a mechanism consisting in loading cohesin complexes at transcribed enhancers or promoters (18–20) followed by chromatin extrusion until cohesin meets convergently oriented CTCF sites (5). In the specific case of *IgH* locus, cohesin is loaded at enhancers (either *3’RR* or *Eμ* regions) to first initiate the “big chromatin loop” between *3’RR* and *Iμ/Sμ* regions at the resting stage. Upon B cell stimulation, new internal loops are shaped to bring together transcribed S regions in order to form the CSR center (CSRC) (15, 16), a structure that favors deletional CSR recombination (17).

The role of *3’RR* and its components, core enhancers and palindromic structure, remains to be determined in the context of CSR center formation. Here, by taking advantage of two mouse models devoid of either the entire *3’RR* (Δ3’RR (6)) or only its palindromic structure (c3’RR model (7)), we evaluated by multiple approaches the role of *3’RR* core enhancers in B cell nuclear organization and *IgH* locus architecture. Interestingly, we showed that the four 3’RR core enhancers are necessary and sufficient to pre-organize resting B cell nucleus to allow deletional CSR recombination at activated stage.

## Materials & Methods

### Mice

8- to 10- weeks old *wt, Δ3’RR, c3’RR* or *Δhs5-7* (kindly gifted by Barbara Birshtein) mice were bred and maintained in SOPF conditions at 21–23°C with a 12-h light/dark cycle. Procedures were reviewed and approved by the Ministère de l’Education Nationale de l’Enseignement Supérieur et Recherche autorisation APAFIS#16151-2018071716292105v3.

### Cell Culture

Splenocytes were collected, non-B cells were removed with mouse B cell kit isolation from stem cell (REF#19854) to collect resting B cells. *In vitro* activated B cells were obtained after three or four days of culture at 1.10^6^ cells/mL in RPMI 1640 medium supplemented with 10% FCS, 2mM Glutamine (Eurobio), 1% Eagle’s Non-essential Amino Acids (Eurobio), 50U/ml of penicillin-streptomycin (Gibco), 1mM sodium pyruvate (Eurobio), 129 μM 2-βmercaptoethanol (Sigma-Aldrich). Cells were activated by adding 1μg/mL lipopolysaccharide (LPS EB ultrapure Invivogen).

### Flow cytometry and Cell Sorting

To sort transitional B cells, B splenocytes were labeled with anti-B220-APC (RA3-6B2 clone, BD 553092) and anti-CD93-BV421 (AA4.1 clone, BD 561990) conjugated antibodies. Transitional (B220+/CD93+) B cells were sorted on ARIA III FACS (BD Biosciences). Plasmablasts were labelled with anti-B220-BV421 (RA3-6B2 clone, BD 562922) and anti-CD138-APC (281-2 clone, BD 558626).

### DNA 3D-FISH

DNA 3D FISH was performed as previously described (21). Briefly, resting and *in vitro* stimulated B cells for three days were dropped onto poly-L-lysine slides and fixed with 4% paraformaldehyde for 10min at room temperature (RT). After PBS washes, cells were permeabilized with pepsin 0.02%/HCl 0.1M for 10min at 37°C. Then cells were washed with PBS and post-fixed with 1% paraformaldehyde for 5min at RT. DNA and probes were denatured in 70% formamide/2X SSC at 72°C and at 95°C for 5min respectively. DNA and probes were hybridized at 37°C overnight. The *IgH* and γ-satellite probes (which highlight heterochromatin) were respectively labelled with dCTP-biotin or dUTP-digoxigenin (Invitrogen REF#19518018 or #11573152910) and dUTP-Alexa Fluor 488 (Invitrogen REF#C11397). Slides were washed in 1X SSC at 72°C for 5min and then incubated with streptavidin-Alexa Fluor 594 (Molecular Probes (life) REF#S32356) (1/300^e^ in 4X SSC) during 1h at RT and mounted with vectashield containing DAPI (Vector Labs). Images were acquired with an epifluorescence microscope (NIKON). Optical sections separated by 0.2μm were captured and stack were deconvoluted and analyzed using Huygens and Volocity softwares, respectively. Separation of alleles was measured from the center of each signal. Volumetric pixel size was 0.065μm in xy and 0.2μm in z-direction.

### ATAC-Seq

10,000 *in vitro* stimulated cells for three days were centrifuged at 500g at 4°C during 10min. Then cells were incubated during 3min in lysis buffer (0,1% Tween-20, 0,1% Nonidet P40, 0,01% Digitonin, 10mM Tris-HCl pH7.4, 10mM NaCl, 3mM MgCl_2_) on ice and resuspended in 10μl of transposition mix (1X TD Buffer from Illumina, Transposase from Illumina, 0,01% digitonin, 0,1% Tween-20). Tagmentation was performed on a thermomixer at 1000rpm 37°C during 30min. Transposed DNA was cleaned-up with minElute Qiagen kit (#28006). Transposed DNA was amplified by PCR using primers Ad1 (5’-AATGATACGGCGACCACCGAGATCTACACTCGTTCGGCAGCGTCAGATGTG-3’) and Ad2.X (5’-CAAGCAGAAGACGGCATACGAGAT**BARECODE**GTCTCGTGGGCTCGGAGATGT-3’). Libraries were sequenced 2*100bp on NovaSeq 6000.

### ATAC-qPCR

DNA accessibility was quantified in resting B cells by qPCR using Bioline SYBR Hi-ROX kit (BIO-92020). Accessibility of Cμ region was quantified using Cμ-Forward (5’-CTTCCCAAATGTCTTCCCCC-3’) and Cμ-Reverse (5’-TGCGAGGTGGCTAGGTACTTG-3’) as well as Cγ1 using Cγ1-Forward (5’-GCCCAAACTAACTCCATGGTC-3’) and Cγ1-Reverse (5’-CAACGTTGCAGGTGACGGT-3’). Cμ and Cγ1 accessibility were normalized to Lig1, using Lig1-Forward (5’-CTCTTCTCCCCGACTGTCAC3’) and Lig1-Reverse (5’-GAGGCTGCTGGGAGTTGTAG-3’), gene identified as invariant between our sample as described in the “analysis of ATAC-Seq section”.

### 3C-HTGTS

3C-HTGTS was performed as previously described (5). Briefly, 10 million of resting or three days stimulated B cells were crosslinked with 2% formaldehyde 10% FCS PBS for 10min at RT under rotation. Crosslinking was stopped by adding glycine at 0.1M. Then, cells were lysed in 50mM Tris, 150mM NaCl, 5mM EDTA, 0.5%NP-40,1% TX-100 supplemented with protease inhibitor (ROCHE #11873580001). Nuclei were resuspended in 0,3% SDS for 1h at 37°C at 900rpm and then neutralized with Triton TX-100 for 1h. DNA restriction was performed using CviQ1 (Thermo Fisher ER0211) in B buffer (Thermofisher #BB5) overnight at 37 °C, before heat inactivation for 25 min at 65 °C. Overnight ligation was performed at 16°C at 300rpm. Next, DNA was treated by proteinase K and RNase and cleaned by phenol/chloroform. After 3C step, the LAM-HTGTS protocol was performed (22). Briefly, 3C DNA was sonicated using the Bioruptor (Diagenode; two pulses at low intensity for 20s), and 10 μg was used for the LAM-HTGTS step. A 3′CBE (5’biotin-CACTGTCCAGACAGCAAACC-3’), Iμ/Sμ (5’biotin-GCAGACCTGGGAATGTATGGT-3’) or 3’RR (5’biotin-GGACTGCTCTGTGCAACAAC-3’) bait was used for primer elongation. These single-stranded DNA fragments were incubated with streptavidin beads (Dynabeads C1 streptavidin beads; Invitrogen) overnight at RT and washed with BW buffer (1M NaCl, 5mM Tris-HCl pH7.4, 0,5mM EDTA pH 8.0). A universal I7 adaptor (5’-GCGACTATAGGGCACGCGTGGNNNNNN-3’NH2 and 5’-P CCACGCGTGCCCTATAGTCGC-3’NH2) was ligated before the nested PCR performed with 3′CBE-bait nested primer (5’-ACACTCTTTCCCTACACGACGCTCTTCCGATCT**BARECODE**ACCGGCATGTTCATCAACAC-3’) or Iμ/Sμ-bait nested primer (5’-ACACTCTTTCCCTACACGACGCTCTTCCGATCT**BARECODE**ACACAAAGACTCTGGACCTC-3’) or 3’RRbait nested primer (5’ ACACTCTTTCCCTACACGACGCTCTTCCGATCT**BARECODE**CAAGCTGGGGTCAGAGCATG-3’) and universal I7 reverse (5’-CTCGGCATTCCTGCTGAACCGCTCTTCCGATCTGACTATAGGGCACGCGTGG-3’). After the Tagged PCR with I7 and I5 Illumina primer, PCR products were cleaned using PCR clean-up kit (Macherey-Nagel REF#740609) and validated after migration on BioAnalyser (Agilent). 3C-HTGTS libraries were sequenced 300bp paired-end MiSeq V3 with 20% PhiX.

### LAM-HTGTS

Genomic DNA (gDNA) was extracted from 10 million of 4 days stimulated B cells. LAM-HTGTS was performed as previously described (22) with a *Sμ* bait. Data analysis of MiSeq sequencing reads was performed as previously described (23). All sequence alignments were done with the mouse mm10 genome. Analysis of junction structure has been performed with CSReport tool (24).

### Analysis of ATAC-seq

Nextera sequences were trimmed by TrimGalore (25) with only pairs of reads with their two reads of at least 55bp each were reported in the resulting fastqs and then processed by the nf-core/atacseq pipeline (26) using mm10 for reference genome. Differentially opened chromatin regions from comparisons of different conditions were extracted using DESeq2 (27) from the *DESeqDataSet* previously produce by the pipeline (26).

The normalized counts extracted from the *DESeqDataSet* were used to calculate the coefficient of variation (CV) of each consensus peaks. Peaks with coverage greater than or equal to the mean were ranked by increasing CV to find a suitable region for ATAC-qPCR normalization controls among the samples compared. The third least variable consensus peak hitting Lig1 gene from positions 13278266 to 13279324 of chromosome 7 on mm10 were selected to design primers for ATAC-qPCR normalization controls. Chromatin A/B compartments were inferred from the consensus peaks of nf-core/atacseq analysis (26) at 3000kb resolution by an extended version (github trichelab/compartmap repository) of the original R package Compartmap (28).

### Analysis of 3C-HTGTS

Sequencing reads were aligned to the mm10 genome and processed as previously described (29). Each 3C-HTGTS library plotted for comparison was normalized by randomly selecting a number of junctions equal to the total number of junctions present in the smallest library in the comparison set.

### Statistical analysis

Mann Whitney two-tailed tests were used for statistical analysis using GraphPad Prism software (*p<0.05, **p<0.01, ***p<0.001, ****p<0.0001).

## Results

### 3’RR core enhancers maintain *IgH* loci in an active compartment in mature resting B cells

To study the role of 3’RR core enhancers *vs* 3’RR palindromic structure, we compared two mouse models carrying total or partial deletion of the 3’RR to *wt* mice. We took advantage of *c3’RR* model (7) carrying only the four core enhancers and *Δ3’RR* model (6) carrying total deletion of the 3’RR region (Figure 1A). We compare, in these models, transitional splenic B cells (T) collected by flow cytometry (Supplementary Figure S1A), mature resting B cells (R) and *in vitro*-activated B cells for 3 days with LPS (S). To avoid any impact of cell subtypes representation in bulk culture at day 3, we verified the % of CD138 positive cell after LPS stimulation in the three models. As expected, the percentage of newly generated plasmablasts *in vitro* in the presence of LPS was comparable (Supplementary Figure S1B). By using ATAC-Seq (Assay for Transposase-Accessible Chromatin) technic, we determined, with a high throughput approach, chromatin accessibility across the genome in resting and *in vitro* activated B cells from *wt, c3’RR* and *Δ3’RR* mice. First of all, a principal component analysis was performed to verify reproducibility between samples (Supplementary Figure S2). Then, we first analyzed all differentially opened (DO) chromatin regions. As expected a large number of DO regions was identified upon B cell stimulation (Supplementary Table S1A), corresponding to the induction of B cell activation program. In contrast, when we compared mutants to *wt* in resting and in *in vitro* stimulated B cells, only few differentially open chromatin regions were identified (less than 80 regions) (Supplementary Tables S1B and S2). Moreover, when we confronted the % of A and B compartments, no drastic change was observed between mutants nor in resting B cell neither in *in vitro* stimulated B cells (Figure 1B). Then, we focused exclusively on the *IgH* locus, we observed a decrease of its accessibility in resting B cells from *Δ3’RR* mutants in comparison to *wt* (Figure 1C, Top). To confirm this trend, we quantified ATAC-seq normalized read count in three distinct regions across the *IgH* locus (R1: around *Eμ/Sμ*, R2: constant regions and R3: *3’CBE*; whose coordinates are available in Supplementary Table S3) and we observed a global decrease of *IgH* accessibility throughout the 3’ part of the locus (Figure 1C, Middle). In parallel, we aimed to confirm this result by quantifying DNA accessibility by qPCR. For this purpose, we first identified an open region with low variability between samples to normalize qPCR. As expected, we noticed a slight decrease of *IgH* accessibility in both constant regions tested in *Δ3’RR* resting B cells (Figure 1C, Bottom). Surprisingly, when we compared *c3’RR* deficient and *wt* mice, we showed that the *c3’RR* resting B cells harbour the same profile as the *wt*, demonstrating that the 3’RR core enhancers are sufficient to maintain *IgH* loci in an open chromatin conformation (Figure 1C). In LPS-stimulated B cells, *IgH* locus is also globally less accessible in *Δ3’RR* model (Figure 1D). A slight decrease of IgH accessibility was shown in the *c3’RR* model across the locus.

Subsequently, we assessed addressing of *IgH* loci to pericentromeric heterochromatin (PCH) within B cells nuclei by a DNA 3D-FISH (Fluorescent *In Situ* Hybridization) approach using an *IgH* fluorescent probe (RP23-109B20) encompassing a region lying from the *D*_*H*_ cluster to *Cγ2a* segment (Figure 2A) and a probe (major γ-satellite) specific to PCH as the “inactive zone”. In a wild-type context, in all three cell types tested, approximately 20% of cells have at least one *IgH* allele in the PCH, with a minor proportion of cells having both *IgH* alleles in PCH (Figure 2B, left). In *Δ3’RR* mice, around 15% of *IgH* alleles are localized in PCH at T stage as in *wt* mice. In contrast, in resting B cells, the proportion of B cells harboring at least one *IgH* allele colocalized to PCH reaches 35% and includes a higher proportion of cells for which both alleles localized to PCH (Figure 2B, middle). This configuration is maintained in *in vitro* stimulated B cells. Surprisingly, when only 3’RR core enhancers, not the surrounding DNA packaging, are reintroduced (*c3’RR* model), almost the same profile as in *wt* was observed (Figure 2B, right). Overall, DNA 3D-FISH and ATAC-seq experiments showed a global decrease of *IgH* loci accessibility in resting B cells and in *in vitro* stimulated B cells from *Δ3’RR*. In contrast, *c3’RR* B cells are in an open chromatin configuration almost as in the *wt* context. All together these results show that in *wt* resting B cells, *IgH* loci were widely in an open chromatin compartment and that PCH addressing is governed only by 3’RR core enhancers, and not by palindromic structure. These findings point out a crucial role of the *3’RR* core enhancers in regulating the nuclear organization of mature resting B cells.

**Figure 2.**
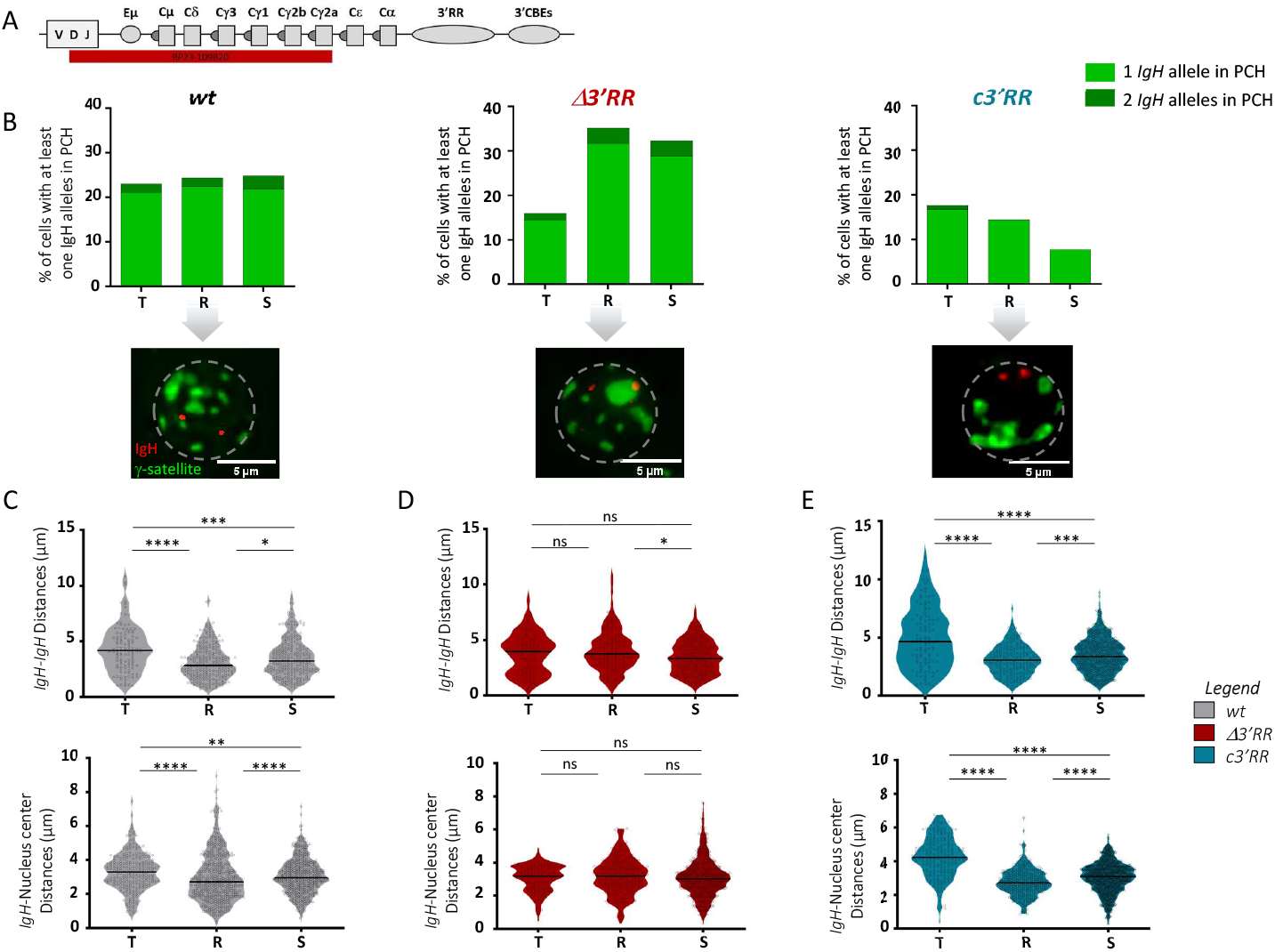
*3’RR* core enhancers are sufficient to maintained *IgH* loci close together and close to the center in mature resting B cells. **A**. Localization of RP23-109B20 FISH probe is represented in red. **B**. Top: Bars graphs represent the % of *IgH* loci into pericentromeric heterochromatin (PCH, stained by γ-satellite major probe) in transitional (T) (*wt* = 116 nuclei analyzed from 3 independent mice) (*Δ3’RR* = 75 nuclei analyzed from 3 independent mice) (*c3’RR* = 108 nuclei analyzed from 2 independent mice), in resting (R) (*wt* = 232 nuclei analyzed from 3 independent mice) (*Δ3’RR* = 136 nuclei analyzed from independent mice) (*c3’RR* = 199 nuclei analyzed from 3 independent mice) and in activated (S) (*wt* = 239 nuclei analyzed from 4 independent mice) (*Δ3’RR* = 249 nuclei analyzed from 4 independent mice) (*c3’RR* = 740 nuclei analyzed from 3 independent mice) B cells by DNA-3D-FISH. Light and dark green respectively corresponds to the % of cells harboring one or two *IgH* allele in PCH. Bottom: Representative nuclei of resting B cells from each model (*IgH* loci in red / PCH in green, scale bars: 5μm). **C**. Violin plots represent raw *IgH-IgH* interallelic distances in μm from transitional to stimulated B cell stages in each model. **D**. As in (**C**) for *IgH*-nucleus center distances.

### 3’RR core enhancers are sufficient to regulate *IgH* loci position dynamics in nuclei of mature B cells

Using DNA 3D-FISH, we also investigated the position of *IgH* loci within nuclei of transitional, mature resting and *in vitro* stimulated B cells from the three mouse models. We measured the *IgH* inter allelic distance as well as the distance between *IgH* loci and nuclear center at each stage of differentiation cited above. In the *wt* context, we observed that during the transition from transitional to mature resting B cell stage, *IgH* loci move closer together (p<0.0001), and then, move away from each other during *in vitro* activation (p=0.0122) (Figure 2C, Top). In *wt* mice, the same dynamic is observed when we measured the distances between *IgH* loci and the nuclear center: during transition from transitional to resting stage, *IgH* loci move closer to the center of nucleus (p<0.0001), while they move away from it upon *in vitro* stimulation (p<0.0001) (Figure 2C, Bottom). In contrast, B cells deficient for the entire 3’RR (*Δ3’RR* model) loss the dynamics observed in *wt* context. In fact, the *IgH* interallelic distance remains constant (Figure 2D, Top). Similarly, distance between *IgH* loci and nucleus center does not differ through transitional to resting and stimulated stages (Figure 2D, Bottom). Interestingly, the reintroduction of *3’RR* core enhancers (*c3’RR* model) alone rescued normal dynamics: *IgH* loci get closer to each other during transitional to resting stage evolution (p<0.0001) and move away from each other upon *in vitro* stimulation (p=0.0004) (Figure 2E, Top). Comparably, in *c3’RR* resting B cells, *IgH* loci are closer to the nucleus center than in transitional B cells (p<0.0001) and then, *IgH* loci move away from nucleus center during activation (p<0.0001) (Figure 2E Bottom). These results pinpoint a dynamic relocalization of *IgH* loci during late B cell maturation orchestrated by the *c3’RR* enhancers alone and highlight the crucial role carried by the 3’RR core enhancers in the nuclear positioning of *IgH* loci. They allow both the dynamic relative positioning of *IgH* loci to each other, and to the center of nucleus, especially at resting B cell stage. So, it is tempting to speculate that the 3’RR core enhancers pre-organize resting B cell nucleus in order to properly position *IgH* loci for remodeling events occurring upon activation.

### 3’RR core enhancers allow *IgH* loop conformation in mature resting and stimulated B cells

As a supplemental level of regulation, chromatin loop formation appears as a master regulator of gene remodeling, such as CSR, in B cells. But, the regulation of loop formation itself is not fully described. Our mouse models allowed to assess the respective involvement of components of the *3’RR* in *IgH* locus looping dynamics. To achieve this point, we performed, in resting and LPS-stimulated B cells from *wt* and mutants, high resolution 3C-HTGTS (15) with two complementary viewpoints: one located within the *Iμ/Sμ* region (named *Iμ/Sμ* bait) (Figure 3A) (17) and another one located between hs5 and hs6 enhancers within the 3’ *IgH* TAD border region conserved between the three models (named *3’CBE* bait) (Supplementary Figure S4A) for quantifying chromatin loop from bait to prey (Supplementary Table S4A for prey coordinates).

**Figure 3.**
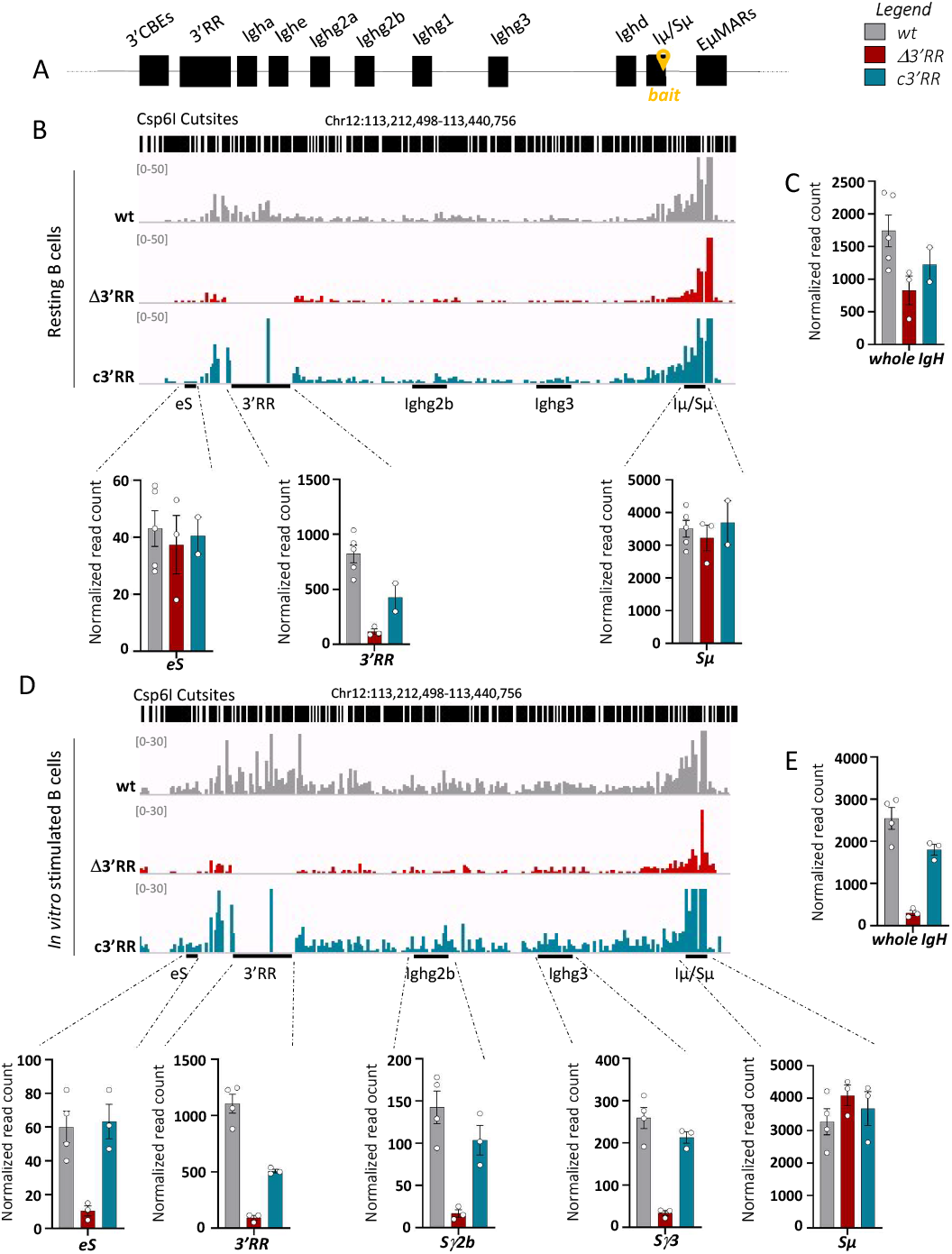
3’RR core enhancers are mandatory for *IgH* big loop and internal loops formation at resting and stimulated B cell stages. **A**. Representation of murine *IgH* locus, the yellow arrow represents the 3C-HTGTS bait. **B**. Top: Representative bedgraphs from splenic resting B cell sample for *wt* (n=5 independent mice), *Δ3’RR* (n=3 independent mice) and *c3’RR* (n=2 independent mice) models, visualized with IGV. Bedgraphs were normalized to smallest sample. Bottom: Bargraphs of normalized reads count. **C**. Quantification of normalized reads count in the entire *IgH* locus in splenic resting B cells from *wt* and mutant mice. **D**. Same as in (**B**) in *wt* (n=4 independent mice), *Δ3’RR* (n=3 independent mice) and *c3’RR* (n=3 independent mice) *in vitro* LPS-stimulated B cells. **E**. Same as in (**C**). Scale is adapted to see interactions of interest.

First, 3C-HTGTS performed in *wt* resting B cells with *Iμ/Sμ* bait showed long-range contacts between *Iμ/Sμ* and the *3’RR* region (Figure 3B bedgraphs and right bargraph, Supplementary Figure S3B and Supplementary Table S4B for individual mice); these data reflect the first level of *IgH* locus folding (“big loop”) and are consistent with both pioneer studies based on chromosome capture (14) and the more recent studies built on high throughput techniques (15). In contrast and as expected, *Δ3’RR* resting B cells were not enable to perform any chromatin loop between *Iμ/Sμ* and the *3’RR* region as a consequence of the total deletion of the *3’RR* region (Figure 3B bedgraphs and middle bargraph and Supplementary Figure S3B). In this model, the decrease of long-range (*3’RR*) interactions from *Iμ* was not compensated by any new interaction inside the locus nor outside the locus at the *eS* region, a phenomenon previously described in the mouse model devoid of the *3’CBE* region (5) (Figures 3B bedgraphs and left bargraph and C). Strikingly, in resting *c3’RR* B cells, devoid of the palindromic structure, chromatin loops between *Iμ/Sμ* and *3’RR* regions still occurred notably thanks to the hs1.2 enhancer (Figure 3B bedgraphs and right bargraph and Supplementary Figure S3B). This finding was confirmed, albeit to lesser extent, by using the *3’CBE* bait located downstream from 3’ super-enhancer (Supplementary figure S4A). Similar chromatin loops were observed in *wt* and *c3’RR* model (Supplementary figure S4B), harboring both high interaction frequencies between the *Iμ/Sμ* and the *3’RR* regions. In contrast, in *Δ3’RR*, 3C-HTGTS did not detect any chromatin loop between *Iμ/Sμ* and the 3’ super-enhancer (Supplementary figure S4B). Altogether, these results highlight, for the first time, a major role of 3’ regulatory enhancers in *IgH* locus folding (“big loop”, involving *Iμ/Sμ* and the 3’ super-enhancer) of resting B cells. While the *3’RR* core enhancers are necessary and sufficient for the *IgH* “big loop” formation, the *3’RR* DNA packaging (supporting the quasi-palindromic organization) is not essential for such structure.

Secondly, we analysed in *in vitro* stimulated *wt* B cells with LPS for 3 days the occurrence of chromatin loop by using the *Iμ/Sμ* viewpoint and globally we found the same pattern as in resting B cells but with a more pronounced difference between genotypes. In *wt*, DNA loop detected by 3C-HTGTS revealed the “big loop” structure and the occurrence of new interactions between *Iμ/Sμ* and the targeted *Sγ3* and *Sγ2b* regions that normally undergoes germline transcription in such conditions (15). In accordance with the literature, this data reflects the physiological *IgH* “internal loops” formed with targeted switch regions in a CSRC (Figure 3D, Supplementary Figure S3C and Supplementary Table S4C for individual mice) (15). In *Δ3’RR* model, the difference is even greater at this stage than in resting B cells, the I*gH* big loop is still not detectable, and a drastic decrease of contact frequencies between *Iμ/Sμ* and *Sγ3* or *Sγ2b* regions is observed (Figure 3D bedgraphs and bargraphs and supplementary figure S3C). Just as in resting B cells, the reduction in long-range interactions upon *3’RR* deletion (Figure 3D) is not compensated by any random interactions within *IgH* locus (Figures 3D bedgraphs and 3E). Interestingly, by using *Iμ/Sμ* bait in LPS-stimulated B cells from *c3’RR* model, the *IgH* big loop is still detected, but slightly decreased because of the lack of palindromic structure. As in resting B cells, interactions from *Iμ/Sμ* to *3’RR* are mainly enabled by hs1.2 enhancer (Figure 3D bedgraphs and right bargraph). On the other hand, contact frequencies between *Iμ/Sμ* and *Sγ3* or *Sγ2b* regions occurred in *c3’RR* model at the same level than in *wt* mice (Figure 3D bedgraphs and bargraphs). In LPS stimulated B cells, 3C-HTGTS performed with the 2^nd^ viewpoint (3’CBE bait) confirmed all these findings. Indeed, high levels of interaction between the bait and *Iμ/Sμ* region were detected in *wt* and *c3’RR* mice as well as internal loops involving the *Sγ3* or *Sγ2b* regions (Supplementary figure S4C). And, as expected, there is still a defect of interactions between *Iμ/Sμ* and targeted switch regions in stimulated *Δ3’RR* B cells (Supplementary Figure S4C).

These data support several hypotheses regarding the establishment of *IgH* loops during B cell activation. Firstly, it seems quite clear that the 3’RR core enhancers alone, without the packaging of surrounding DNA, are sufficient for the formation of *IgH* chromatin loops, the “big loop” and the internal loops, in resting and in activated B cells. Second, the big loop formation at resting stage is a prerequisite for the formation of the internal loops and the CSRC formation in activated B cell. The internal loop defect identified by 3C-HTGTS, reflect the failure to form the CSRC which is in accordance with the strong CSR defect observed in the *Δ3’RR* model (30)

To fully conclude on the role of *3’RR* core enhancers as *IgH* chromatin organizers, we performed 3C-HTGTS on a mice model carrying the partial deletion of *3’CBE* region (deletion from hs5 to hs7) (31) with a bait conserved upon deletion of the entire *3’RR* localized downstream *hs4* (bait 3’RR) (Supplementary Figures S5 A and B). In this model, we observe the formation of the *IgH* big loop in resting B cells as well the internal loops in stimulated B cells. All together these results showed that the *3’CBE* region is not involved in *IgH* chromatin loops formation (Supplementary Figures S5 C and D).

### 3’RR core enhancers are required for proper deletional CSR mechanism

Productive CSR events are the result of deletional recombination events that occur by joining the correctly juxtaposed *Sμ* donor and *Sx* acceptor regions in such a way as to achieve deletion of intermediate DNA sequences. In some cases, the recombination event breaks and religates the large intermediate DNA segment between the two S regions in an inverted manner, resulting in an inversion of the *IgH C* gene, hence called inversional CSR, thereby prohibiting functional expression of heavy chain (Figure 4A). To further investigate the involvement of *3’RR* core enhancers in CSR center formation, we questioned the CSR mechanism involved (*ie*: CSR deletional *vs* CSR inversional) in our mutants. After *in vitro* activation with LPS for four days, we first compared the occurrence of CSR junctions from *Sμ* to *Sγ3* and *Sγ2b* by LAM-HTGTS using a bait localized within *Sμ* (Figure 4B) as previously described (19, 23) and the resulting sequences have been analyzed through the dedicated pipeline. As previously described, the frequency of CSR events in the *c3’RR* model is almost comparable to the *wt* (7) (Figure 4C). Despite the drastic decrease of CSR characterizing the *3’RR*-deficient model (6), few CSR junctions could thus be detected in LPS-stimulated cells (Figure 4C). To identify the CSR mechanism involved in our mutants we analyzed the orientation of the S acceptor region (*i*.*e* +1 or -1). As expected, in the *wt* context, almost all CSR junctions were typical to the CSR deletional mechanism (Figure 4D). In accordance with internal *IgH* loop formation in the *c3’RR* model, all junctions were in favor of deletional CSR (Figure 4D). At the opposite, all the junctions identified in the context of the *3’RR* deletion were characteristic of the inversional CSR mechanism. We then analyzed structural features of CSR junctions and found that the large majority of junctions are blunt, in all three models, a typical hallmark of the NHEJ repair pathway (Figure 4E). Increase of inversional CSR were reported in case of specific DNA repair actors inhibition/deletion (33), however our findings suggest inversional CSR find in our *Δ3’RR* mutants is related to the impairment of loops and not to defect of DNA repair actor recruitment. Indeed, the junctions were almost all blunt suggesting a normal DNA repair pathway (Figure 4F). All together this result suggested that the chromatin loops at the *IgH* locus allow productive CSR recombination by preventing inversional CSR.

**Figure 4.**
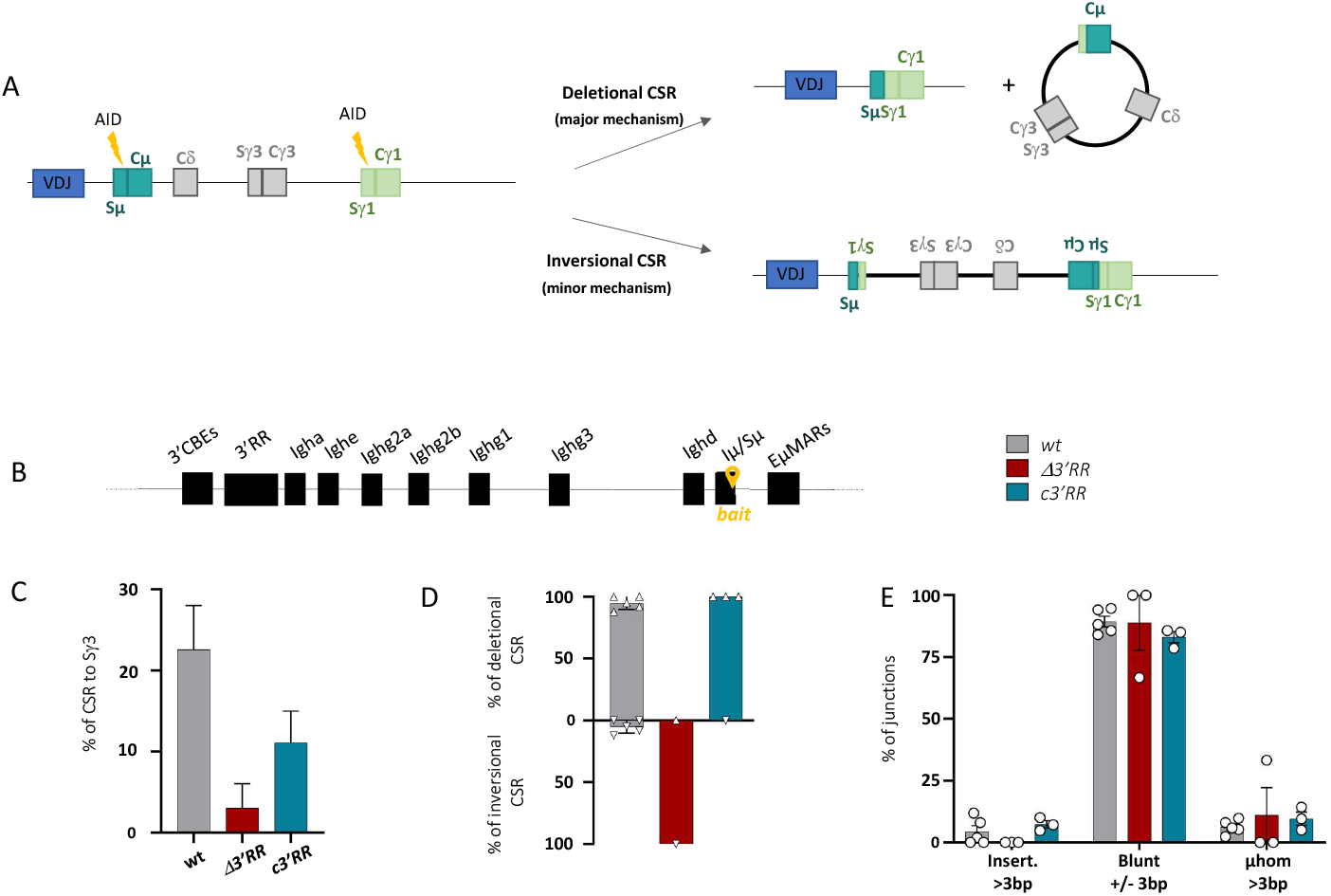
Deletional CSR in stimulated B cells requires 3’RR core enhancers. **A**. Schematic representation of deletional and inversional CSR mechanisms **B**. Localization of *Sμ* bait used is represented by the yellow arrow in the *IgH* locus. **C**. Percentage of switch recombination to Sγ3 determined by LAM-HTGTS in *wt* (n=5 independent mice), *Δ3’RR* (n=3 independent mice) and *c3’RR* (n=3 independent mice) from *in vitro* stimulated B cells. **D**. Proportion of deletional and inversional CSR determined by LAM-HTGTS, in *in vitro* stimulated B cells from *wt* and mutants’ mice. **E**. Structure of CSR junctions analyzed with CSReport.

## Conclusion

Our study clearly demonstrates that 3’RR core enhancers are necessary and sufficient to preorganize the resting B cell nucleus. Firstly, we show that they are mandatory for maintaining *IgH* loci in a transcriptionally active compartment. Secondly, we demonstrate that core enhancers keep *IgH* loci close to each other and close to the nucleus center in an active compartment at resting B cell stage. Finally, we prove that they are essential for forming the *IgH* big loop between *Iμ/Sμ* and the *3’RR* regions prior the internal loops involving the *S* regions at activated B cells stage. Altogether our results highlight that *3’RR* core enhancers, beyond their regulatory function on *IgH* transcription (32), regulate *IgH* chromatin loops formation to prepare *IgH* locus to CSR. We propose a simplistic model in which *3’RR* core enhancers shape, by themselves, resting B cells at nuclear and supranucleosomal levels (See graphical abstract).

As previously described, in *wt* resting B cells, a small proportion of cells harbors an *IgH* locus localized in pericentromeric heterochromatin (35, 36). However, in *in vitro* stimulated B cells, their relative positioning is still debate. According to Skok’s lab, the unproductive *IgH* allele colocalized with PCH and replicated later (35), implying that the non-productive *IgH* allele is excluded only by its nuclear location. In contrast, we and the De Latt group showed, in *in vitro* activated B cells, that both *IgH* loci remain in euchromatin (36). This finding is in accordance with *IgH* biallelic expression observed in mature B cells (37, 38). In this study, we found that *IgH* locus from *Δ3’RR* B cells is less accessible, which is consistent with previous ChIP-qPCR experiments that demonstrated a decrease of active chromatin marks (using anti-H3K9Ac antibody) in both constant and *Sμ* regions in *Δ3’RR* activated B cells (30). This observation perfectly correlates with the BCR low phenotype found in this model in mature B cell subpopulations and enforces the role of *3’RR* in late stages of B cell differentiation (39). Moreover, *IgH* locus accessibility is in agreement with *IgH* constant region transcription; in *c3’RR* model the level of constant regions transcription is similar to the *wt* context whereas in *Δ3’RR* mice the transcription is largely decreased but not abolished (7). The reduced accessibility of the *IgH* locus in the *Δ3’RR* model may be explained by fixation of chromatin-regulating factors in the *3’RR*. In particular, the *3’RR* super-enhancer recruits Brg1, a protein forming part of the SWI/SNF chromatin remodeling complex, known to preferentially bind the enhancers with the aim to open the chromatin and activate the transcription. This factor was initially described to bind the *3’RR* in immature B cells (40, 41). More recently, Brg1 was found to bind the 3’RR in mature B cells especially during the GC formation (42). ZMYND8, a protein described as a “chromatin reader”, is also known to bind the *3’RR*. The conditional deletion of ZMYND8 in B cell results in a phenotype similar to the 3’RR deletion (reduced CSR and SHM) suggesting the involvement of this factor in regulation of *IgH* locus at supranucleosomal scale (43).

On another scale, our data show that *3’RR* core enhancers are required for the establishment of chromatin loops at the *IgH* locus during CSR. However the Pavri Group recently published that the *IgH* big loop was mediated only by the *3’CBE* region and not by the *3’RR* region (44). This study was carried out only under LPS+IL4 stimulation conditions, inducing a switch to IgG1, an isotype known to be slightly differently regulated from other isotypes, since the switch to IgG1 persists, to a lesser extent, in the *Δ3’RR* model. Moreover, in the *Δhs567* mice (partial *3’CBE* deletion), it was shown that the *IgH* loops occurred as in a *wt* context in line with a normal CSR recombination (31). In addition, in a model of *3’CBE* total deletion, a decrease of CSR is observed to all isotypes, except IgG1, suggesting a particular link between the *3’CBE* region and the CSR toward IgG1.

*Δ’3RR* and *c3’RR* models showed a drastic decrease in SHM (21) suggesting that the mechanisms governing the nucleus organization and three-dimensional topology of *IgH* locus are different between GC and in *in vitro*-activated B cells. So “big loop”, putting the *EμMARs* and *3’RR* regions in contact, seems to not be involved in regulating the SHM process. This suggests a role of the palindromic structure in the formation of another loop, bringing the *3’RR* and *EμMARs* regions into contact with the rearranged *pV*_*H*_.

Described in 2019 by F. Alt’s team, chromatin loop formation at *IgH* locus is mediated by the loop extrusion mechanism (15). It involves loading of the cohesin complex onto DNA by the NIPBL protein, extrusion of DNA into the ring of cohesin complex until encountering two converging CTCF binding sites, followed by dissociation of the cohesin complex from DNA by the WAPL protein. In the specific case of *IgH* locus, cohesin is loaded at enhancers but in the absence of the entire *3’RR*, no loop was found throughout the IgH locus. It is therefore tempting to speculate that the cohesin complex loading onto the *IgH* locus requires the *3’RR* core enhancers.

## Data Availability

Raw data from ATAC-seq, 3C-HTGTS and LAM-HTGTS have been deposited in the European Nucleotide Archive database under access number PRJEB52320.

## Acknowledgments

We thank BISCEm and the animal core facility team for help with mouse work on both practical and regulatory aspects. We are grateful to Emeline Lhuillier and the Genotoul Plateau GeT-Santé facility () for technical assistance with ATAC-Seq sequencing. We thank Vonick Sibut for bio-informatic analysis advices on ATAC-Seq. We thank Adam Yongxin Ye and Xuefei Zhang for precious advices on 3C-HTGTS experiments (technical part and bio-informatic analysis), Mehdi Alizadeh and Emilie Guérin for advices on sequencing step. We are grateful to Dr Brice Laffleur for fruitfull scientific discussion and critical reading of the manuscript. We also thank Pedro P Rocha for scientific and technical discussions. We thank Christelle Oblet for her invaluable help in creating the graphical abstracts. CB was supported by PhD fellowships from the French Ministère de l’Enseignement Supérieur, de la Recherche et de l’Innovation. MT was supported by PhD fellowships from the French Ministère de l’Enseignement Supérieur, de la Recherche et de l’Innovation and the Fonds Européen de Développement Régional (FEDER). JP was supported by Institut CARNOT CALYM. This work was supported Agence Nationale de la recherche (ANR-21-CE15-0001-0), La Ligue Contre le Cancer (comités 87, 23 to EP and SLN), the Fondation ARC pour la recherche sur le cancer (PhD continuation fellowship to CB and MT).

## Author contribution

CB, MT and SLN performed experiments. EP and SLN conceived and supervised the study. JP performed the bio-informatic analysis. D.R and Z.B were very helpful respectively for ATAC-seq and 3C-HTGTS experiments. CB, EP and SLN wrote the manuscript.

### Competing financial interest

The authors declare no competing financial interests.

---

**Supplementary Figure S1**: **A**. Gating strategy to sort transitional B cells from spleen. **B**. Top: gating strategy to identified plasmablasts *in vitro* stimulated splenic B cells from *wt*. Bottom: Bargraphs represented the % of plasmablast in *wt, Δ3’RR* and *c3’RR* mice (n=6-9).

**Supplementary Figure S2**: Principal Component Analysis of ATAC-Seq of resting (R, n=2) and *in vitro* LPS stimulated (J3, n=1-2) B cells from *wt, Δ3’RR* and *c3’RR* mice.

**Supplementary Figure S3**: **A**. Scheme of murine *IgH* locus. Iμ/Sμ bait is represented by yellow arrow. **B**. Bedgraphs, visualized with IGV, of replicates of splenic resting B cells from *wt* (n=5 independent mice), *Δ3’RR* (n=3 independent mice) and *c3’RR* (n=2 independent mice). **C**. Same as in B for *in vitro* stimulated B cells from *wt* (n=4 independent mice), *D3’RR* (n=3 independent mice) and *c’3RR* (n=3 independent mice) mice. Scale is adapted to see interactions of interest.

**Supplementary Figure S4**: **A**. Scheme of murine *IgH* locus. *3’CBE* bait is represented by yellow arrow. **B**. Bedgraphs, visualized with IGV, of replicates of splenic resting B cells from *wt* (n=2 independent mice), *Δ3’RR* (n=2 independent mice) and *c3’RR* (n=2 independent mice). **C**. Same as in **B** for *in vitro* stimulated B cells from *wt* (n=3 independent mice), *Δ3’RR* (n=2 independent mice) and *c’3RR* (n=2 independent mice) mice. Scale is adapted to see interactions of interest.

**Supplementary Figure S5**: **A**. Scheme of murine *IgH* locus and representation of *Δ3’RR and Δhs567* mouse models. Deletion are represented by red crosses **B**. 3’RR bait is represented by yellow arrow. **C**. Bedgraphs, visualized with IGV, of replicates of splenic resting B cells from *wt* (n=3 independent mice), *Δ3’RR* (n=3 independent mice) and *Δhs567* (n=3 independent mice). **D**. Same as in **C** for *in vitro* stimulated B cells from *wt* (n=3 independent mice), *Δ3’RR* (n=2 independent mice) and *Δhs567* (n=2 independent mice) mice. Scale is adapted to see interactions of interest.

## Notes

### Competing Interest Statement

The authors have declared no competing interest.

