## Supplementary files for "Core enhancers of the 3’RR optimize *IgH* nuclear position and loop conformation for oriented CSR"

### Figure S1

**A**

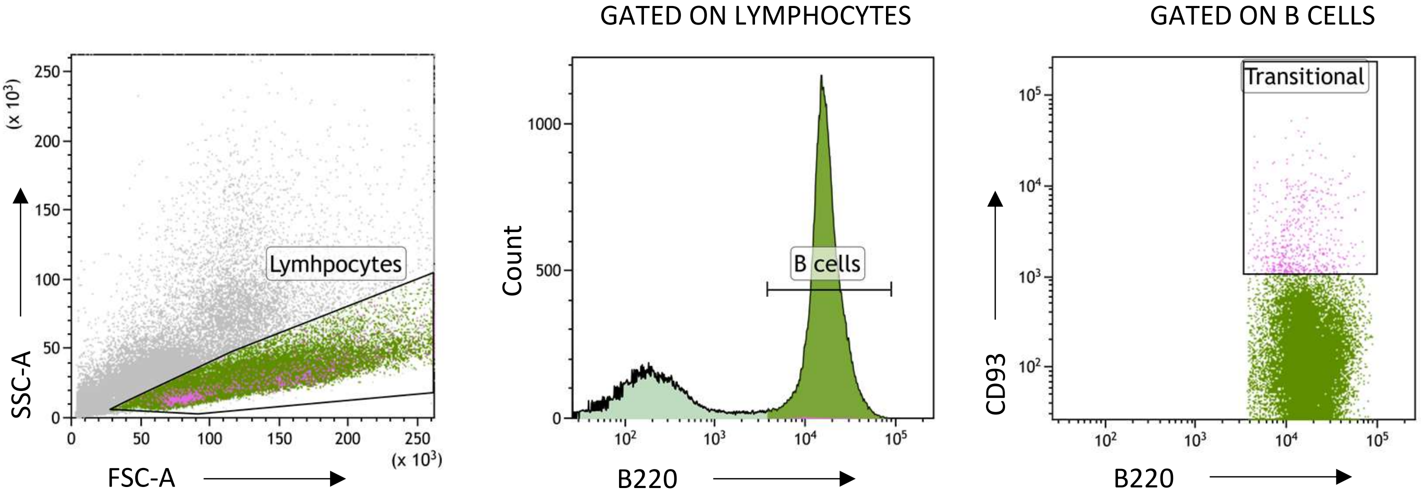


**B**

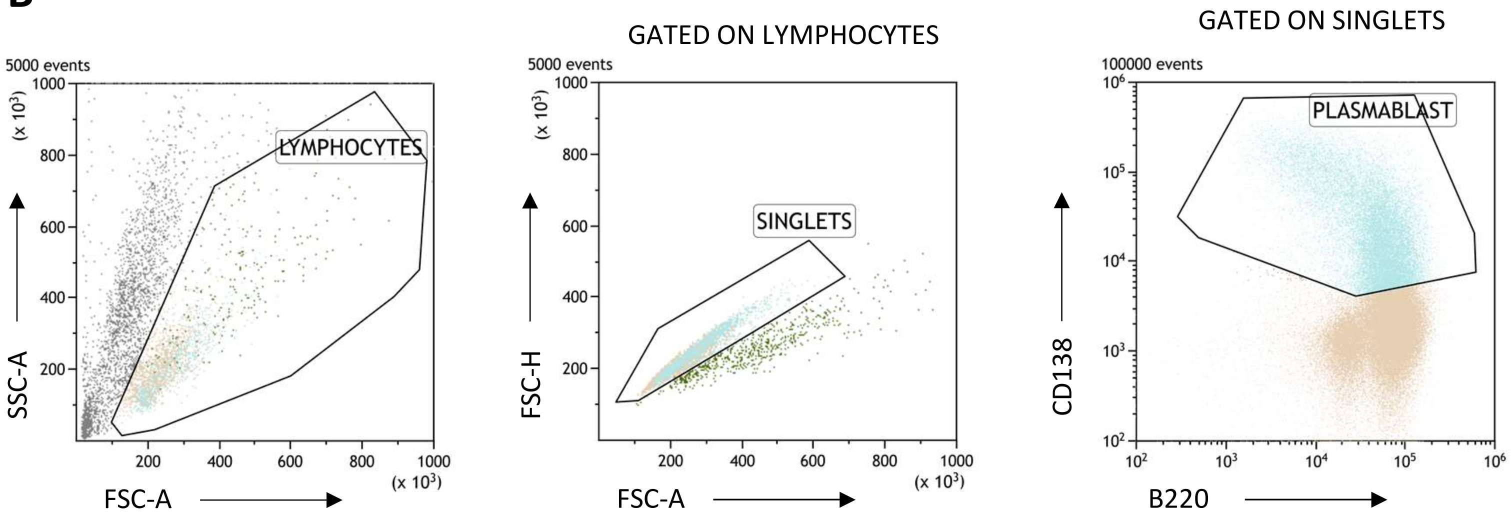


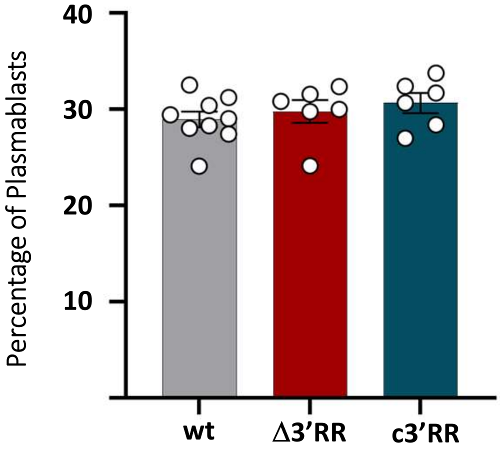


| | wt | Δ3'RR | c3'RR |
|---|---|---|---|
| Mean | 29,0 | 29,8 | 30,7 |

#### Figure S2

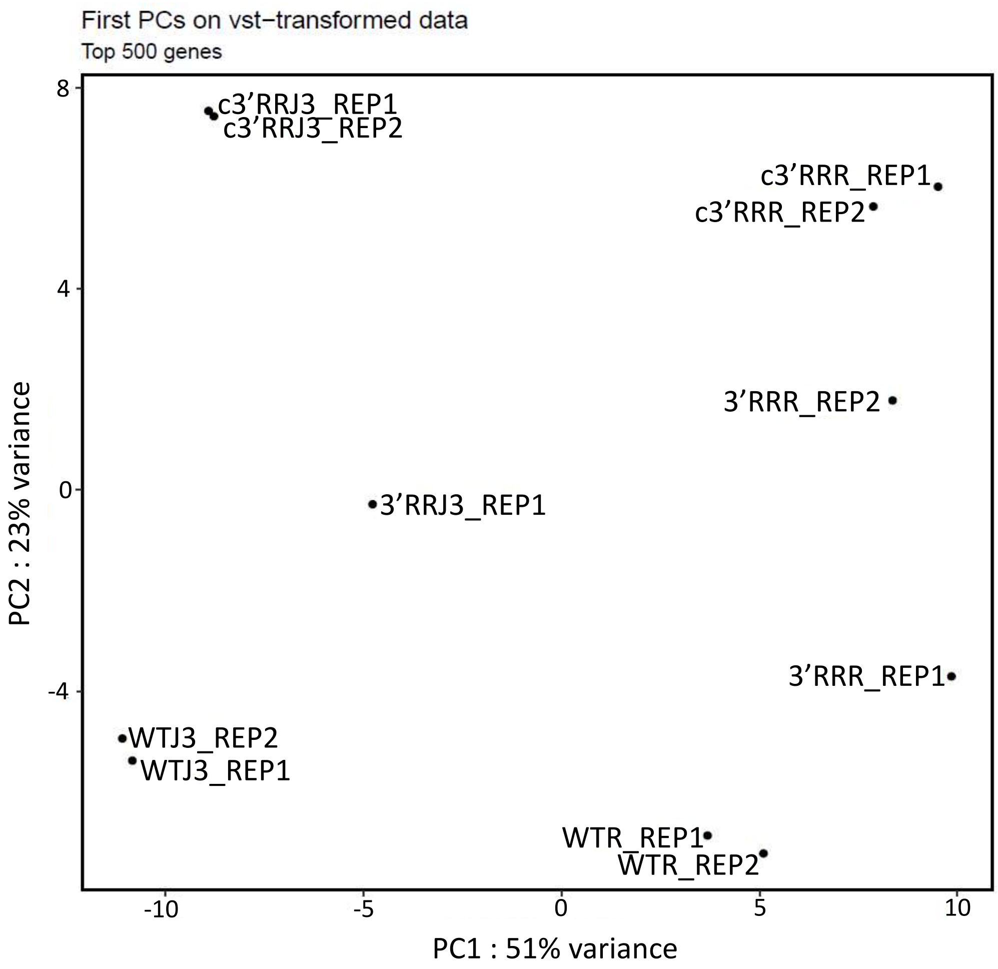
First PCs on vst−transformed data
Top 500 genes
PC2 : 23% variance
PC1 : 51% variance
8
4
0
-4
-10
-5
0
5
10
c3’RRJ3_REP1
c3’RRJ3_REP2
c3’RRR_REP1
c3’RRR_REP2
3’RRR_REP2
3’RRJ3_REP1
3’RRR_REP1
WTJ3_REP2
WTJ3_REP1
WTR_REP1
WTR_REP2

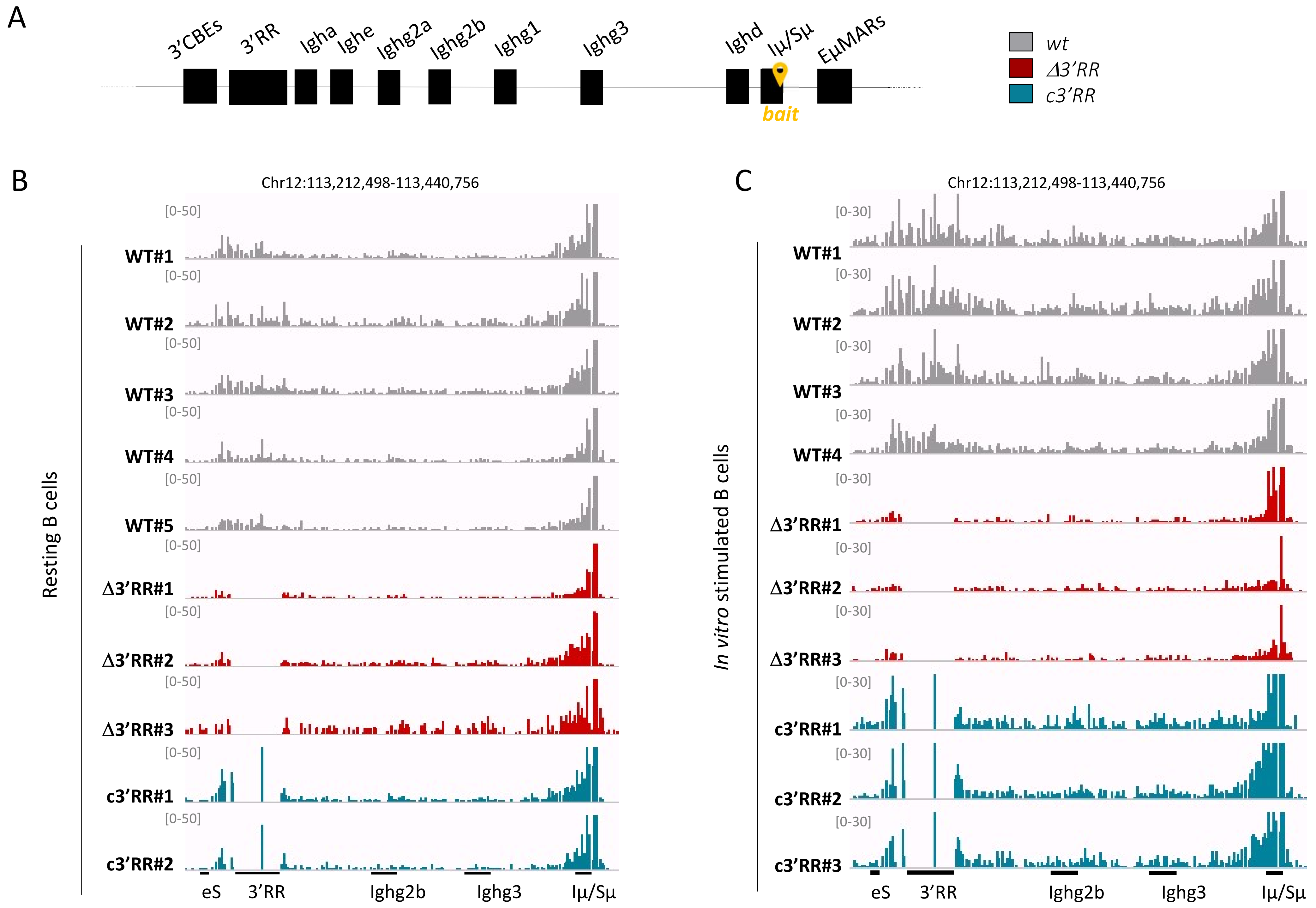
A
3'CBEs
3'RR
Igha
Ighe
Ighg2a
Ighg2b
Ighg1
Ighg3
Ighd
Iμ/Sμ
EμMARs
bait
wt
Δ3'RR
c3'RR
B
Chr12:113,212,498-113,440,756
Resting B cells
[0-50]
WT#1
WT#2
WT#3
WT#4
WT#5
Δ3'RR#1
Δ3'RR#2
Δ3'RR#3
c3'RR#1
c3'RR#2
eS
3'RR
Ighg2b
Ighg3
Iμ/Sμ
C
Chr12:113,212,498-113,440,756
In vitro stimulated B cells
[0-30]
WT#1
WT#2
WT#3
WT#4
Δ3'RR#1
Δ3'RR#2
Δ3'RR#3
c3'RR#1
c3'RR#2
c3'RR#3
eS
3'RR
Ighg2b
Ighg3
Iμ/Sμ

**Figure S3**

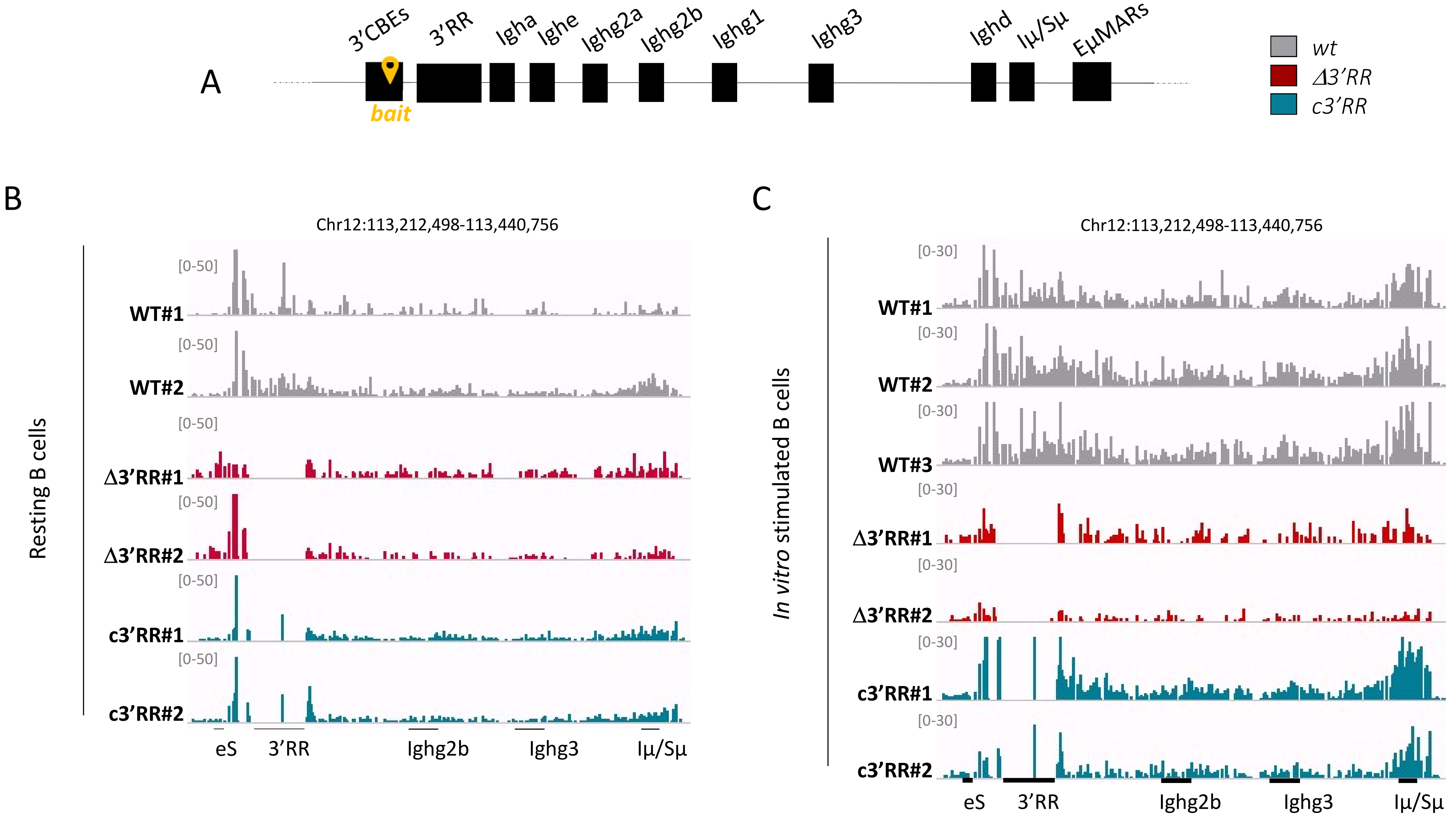
A
3'CBEs
3'RR
Igha
Ighe
Ighg2a
Ighg2b
Ighg1
Ighg3
Ighd
Iμ/Sμ
EμMARs
bait
wt
Δ3'RR
c3'RR
B
Chr12:113,212,498-113,440,756
Resting B cells
[0-50]
WT#1
WT#2
Δ3'RR#1
Δ3'RR#2
c3'RR#1
c3'RR#2
eS
3'RR
Ighg2b
Ighg3
Iμ/Sμ
C
Chr12:113,212,498-113,440,756
In vitro stimulated B cells
[0-30]
WT#1
WT#2
WT#3
Δ3'RR#1
Δ3'RR#2
c3'RR#1
c3'RR#2
eS
3'RR
Ighg2b
Ighg3
Iμ/Sμ

**Figure S4**

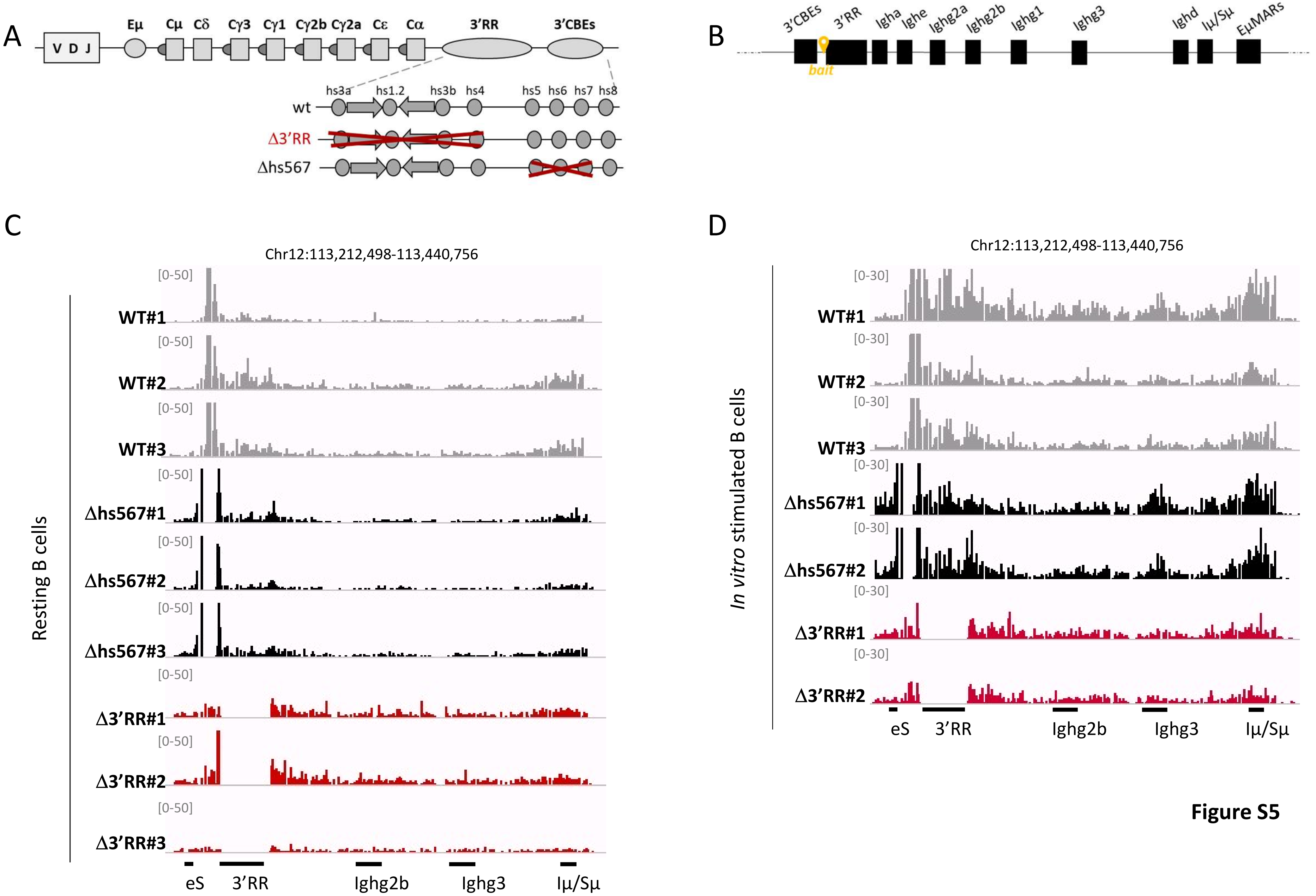
A
V D J
Eμ
Cμ
Cδ
Cγ3
Cγ1
Cγ2b
Cγ2a
Cε
Cα
3'RR
3'CBEs
hs3a
hs1.2
hs3b
hs4
hs5
hs6
hs7
hs8
wt
Δ3'RR
Δhs567
B
3'CBEs
3'RR
Igha
Ighe
Ighg2a
Ighg2b
Ighg1
Ighg3
Ighd
Iμ/Sμ
EμMARs
bait
C
Chr12:113,212,498-113,440,756
Resting B cells
[0-50]
WT#1
WT#2
WT#3
Δhs567#1
Δhs567#2
Δhs567#3
Δ3'RR#1
Δ3'RR#2
Δ3'RR#3
eS
3'RR
Ighg2b
Ighg3
Iμ/Sμ
D
Chr12:113,212,498-113,440,756
In vitro stimulated B cells
[0-30]
WT#1
WT#2
WT#3
Δhs567#1
Δhs567#2
Δ3'RR#1
Δ3'RR#2
eS
3'RR
Ighg2b
Ighg3
Iμ/Sμ

**Figure S5**

#### Table S1 : Number of differentially opened regions by ATAC-Seq

# A

| | Number of differentially opened regions |
|---|---|
| WT R *vs* WT S | 687 |
| c3'RR R *vs* c3'RR S | 1292 |
| 3'RR R *vs* 3'RR S | 384 |

# B

| | Number of Differentially opened regions | |
|---|---|---|
| | **Resting** | ***In vitro* stimulated** |
| WT *vs* c3'RR | 45 | 73 |
| WT *vs* Δ3'RR | 12 | 8 |
| c3'RR *vs* Δ3'RR | 11 | 1 |

#### Table S2 : Differentialy opened regions, compared to *wt* , in resting and stimulated B cells from *Δ3'RR* and *c3'RR* mutant mice

# A

WT *vs* Δ3'RR in resting B cells

| PeakID | baseMean | log2FoldChange | pvalue | Chr | Start | End | Strand | Annotation | Distance to TSS | Gene Name |
|---|---|---|---|---|---|---|---|---|---|---|
| Interval_12080 | 65,32400803 | 1,382533223 | 1,79903E-05 | 11 | 118218906 | 118219606 | + | promoter-TSS (ENSMUSG00000017132) | -6514 | Cyth1 |
| Interval_12599 | 53,35748369 | 2,491795186 | 8,17632E-09 | 12 | 20920526 | 20921709 | + | promoter-TSS (ENSMUSG00000085642) | 145 | Zfp125 |
| Interval_12753 | 33,5991101 | 1,760311902 | 3,70619E-05 | 12 | 31973306 | 31973947 | + | intron (ENSMUSG00000002997, intron 5 of 10) | -33219 | Hbp1 |
| Interval_15943 | 149,6198133 | 0,8875904 | 5,6026E-05 | 13 | 43764957 | 43765971 | + | Intergenic | -19721 | Cd83 |
| Interval_30667 | 55,30642524 | -1,652147387 | 2,11986E-05 | 19 | 21912947 | 21914145 | + | Intergenic | -23834 | Ldhb-ps |
| Interval_31323 | 44,29469979 | -2,069512985 | 4,66191E-07 | 19 | 44351580 | 44352475 | + | Intergenic | 18935 | Scd4 |
| Interval_44362 | 27,05318006 | 2,289530272 | 3,88176E-05 | 5 | 23718896 | 23719554 | + | Intergenic | -6559 | AI506816 |
| Interval_46728 | 14,33030532 | 2,908161161 | 6,42419E-05 | 5 | 122602245 | 122602675 | + | intron (ENSMUSG00000029469, intron 6 of 6) | 4909 | Ift81 |
| Interval_47330 | 53,93385657 | -1,995496625 | 5,65951E-07 | 5 | 139780930 | 139782044 | + | Intergenic | -5849 | Ints1 |
| Interval_47620 | 28,93190946 | 2,987157693 | 7,11953E-09 | 5 | 147049813 | 147050666 | + | intron (ENSMUSG00000016520, intron 1 of 9) | -27325 | Polr1d |
| Interval_56539 | 212,1635314 | 0,844766457 | 1,76651E-05 | 8 | 83739935 | 83742697 | + | promoter-TSS (ENSMUSG00000002885) | 10 | Adgre5 |
| Interval_59014 | 47,91307206 | 1,827564692 | 7,51789E-05 | 9 | 58510246 | 58511060 | + | Intergenic | 21691 | Insyn1 |

# B

WT *vs* c3'RR in resting B cells

| PeakID | baseMean | log2FoldChange | pvalue | Chr | Start | End | Strand | Annotation | Distance to TSS | Gene Name |
|---|---|---|---|---|---|---|---|---|---|---|
| Interval_21965 | 32,17992055 | 3,865821078 | 1,29912E-10 | 15 | 79900715 | 79901440 | + | TTS (ENSMUSG00000009585) | 735 | Apobec3 |
| Interval_56539 | 212,1635314 | 1,18583272 | 7,49072E-10 | 8 | 83739935 | 83742697 | + | promoter-TSS (ENSMUSG00000002885) | 10 | Adgre5 |
| Interval_31323 | 44,29469979 | -2,497613172 | 1,63031E-09 | 19 | 44351580 | 44352475 | + | Intergenic | 18935 | Scd4 |
| Interval_59014 | 47,91307206 | 2,688054574 | 1,66363E-09 | 9 | 58510246 | 58511060 | + | Intergenic | 21691 | Insyn1 |
| Interval_31829 | 42,87674911 | 2,497945969 | 1,99371E-08 | 19 | 61151514 | 61152486 | + | Intergenic | -11183 | Zfp950 |
| Interval_26229 | 208,4196308 | 2,089885175 | 3,08772E-08 | 17 | 30611699 | 30612947 | + | intron (ENSMUSG00000024026, intron 1 of 2) | 246 | Glo1 |
| Interval_22720 | 21,47947807 | 4,10409096 | 1,99232E-07 | 15 | 100443572 | 100443968 | + | Intergenic | -5914 | n-R5s43 |
| Interval_26244 | 38,10047802 | 2,071492998 | 1,14397E-06 | 17 | 30901814 | 30902460 | + | intron (ENSMUSG00000024027, intron 1 of 13) | 278 | Glp1r |
| Interval_8283 | 30,33147907 | -3,054353917 | 1,64249E-06 | 11 | 6497445 | 6498063 | + | Intergenic | -21837 | Purb |
| Interval_150 | 26,71177382 | 2,655475369 | 2,6277E-06 | 1 | 13593256 | 13593668 | + | Intergenic | -3587 | Tram1 |
| Interval_26649 | 137,2364225 | 1,04371443 | 3,53146E-06 | 17 | 36110320 | 36111975 | + | promoter-TSS (ENSMUSG00000092365) | -279 | BC023719 |
| Interval_8250 | 13,00628344 | -7,12937049 | 4,306E-06 | 11 | 6006357 | 6006741 | + | intron (ENSMUSG00000057897, intron 3 of 10) | -22411 | Camk2b |
| Interval_36363 | 22,79411746 | -2,390389571 | 4,99884E-06 | 2 | 173246679 | 173247222 | + | intron (ENSMUSG00000038400, intron 1 of 3) | -28053 | Zbp1 |
| Interval_21963 | 110,4203818 | 1,252723517 | 5,92411E-06 | 15 | 79891525 | 79893135 | + | promoter-TSS (ENSMUSG00000009585) | 2798 | AC113595.1 |
| Interval_12599 | 53,35748369 | 1,940154387 | 8,37645E-06 | 12 | 20920526 | 20921709 | + | promoter-TSS (ENSMUSG00000085642) | 145 | Zfp125 |

| | | | | | | | | | | |
|---|---|---|---|---|---|---|---|---|---|---|
| Interval_23703 | 28,86762533 | 2,160736956 | 1,1068E-05 | 16 | 31066693 | 31067108 | + | intron (ENSMUSG00000047434, intron 1 of 3) | 2926 | Xxylt1 |
| Interval_26645 | 96,3291297 | 1,123886763 | 1,31793E-05 | 17 | 36082985 | 36084629 | + | exon (ENSMUSG00000073406, exon 2 of 6) | 416 | H2-Bl |
| Interval_22557 | 63,60686106 | 1,481887312 | 1,39924E-05 | 15 | 97458206 | 97459223 | + | Intergenic | 97512 | Pced1b |
| Interval_3614 | 11,9978808 | 6,753681783 | 1,75376E-05 | 1 | 171065212 | 171065430 | + | promoter-TSS (ENSMUSG00000059498) | -669 | Mir6546 |
| Interval_49487 | 37,1492157 | -2,247784038 | 3,50483E-05 | 6 | 87457375 | 87457932 | + | TTS (ENSMUSG00000030047) | 5994 | Arhgap25 |
| Interval_60872 | 120,6465451 | -3,752072151 | 3,59424E-05 | 9 | 123936024 | 123937162 | + | Intergenic | 32099 | Ccr1 |
| Interval_36364 | 36,85784676 | -1,867822973 | 5,78303E-05 | 2 | 173263590 | 173264142 | + | intron (ENSMUSG00000038400, intron 1 of 3) | 12323 | Pmepa1 |
| Interval_11216 | 97,55991131 | 1,008032889 | 6,01264E-05 | 11 | 99162626 | 99163659 | + | Intergenic | -8065 | Ccr7 |
| Interval_57274 | 25,84206401 | 1,916226576 | 6,08076E-05 | 8 | 114801723 | 114802128 | + | intron (ENSMUSG00000004637, intron 8 of 8) | 89758 | Wwox |
| Interval_21740 | 187,9637624 | -0,804022157 | 6,61994E-05 | 15 | 76195518 | 76197057 | + | promoter-TSS (ENSMUSG00000022565) | 10034 | Plec |
| Interval_3570 | 180,1793892 | 0,858393433 | 7,82664E-05 | 1 | 170174174 | 170175521 | + | promoter-TSS (ENSMUSG00000026670) | -81 | Uap1 |
| Interval_3802 | 13,31646395 | 3,454270326 | 8,95444E-05 | 1 | 173905654 | 173906189 | + | intron (ENSMUSG00000026536, intron 3 of 6) | 7125 | Ifi211 |
| Interval_21964 | 51,86372983 | 4,339140464 | 9,9423E-05 | 15 | 79894946 | 79896164 | + | exon (ENSMUSG00000009585, exon 2 of 8) | -4787 | Apobec3 |
| Interval_26299 | 51,59054898 | 1,459141889 | 0,000106059 | 17 | 31840333 | 31840838 | + | Intergenic | 10990 | Sik1 |
| Interval_34659 | 63,48790214 | 1,286894861 | 0,000106301 | 2 | 119015023 | 119016216 | + | Intergenic | 7410 | Uqcrh-ps2 |
| Interval_47825 | 71,08511526 | -1,248560772 | 0,000106463 | 6 | 3399079 | 3399916 | + | promoter-TSS (ENSMUSG00000047735) | -14491 | Samd9l |
| Interval_55587 | 22,88391855 | 2,573328868 | 0,000116775 | 8 | 35123345 | 35123955 | + | Intergenic | 9882 | Rpl31-ps23 |
| Interval_31117 | 77,06605739 | 1,142616199 | 0,000120654 | 19 | 38025756 | 38027125 | + | intron (ENSMUSG00000048612, intron 1 of 9) | 16959 | Myof |
| Interval_60871 | 127,8743705 | -2,985733378 | 0,000124221 | 9 | 123930334 | 123930943 | + | Intergenic | 38054 | Ccr1 |
| Interval_3914 | 36,31498783 | -1,593335152 | 0,000140264 | 1 | 177984119 | 177984575 | + | intron (ENSMUSG00000091476, intron 1 of 19) | -7089 | Catspere2 |
| Interval_23653 | 14,59206643 | 2,911240379 | 0,000144414 | 16 | 30155064 | 30155415 | + | Intergenic | -13294 | n-R5s32 |
| Interval_376 | 20,53100598 | 2,382695576 | 0,000178836 | 1 | 33883063 | 33883748 | + | promoter-TSS (ENSMUSG00000042182) | 75 | Bend6 |
| Interval_44362 | 27,05318006 | 2,064182212 | 0,000195559 | 5 | 23718896 | 23719554 | + | Intergenic | -6559 | AI506816 |
| Interval_45629 | 165,5957232 | 0,750055988 | 0,000201062 | 5 | 86065126 | 86066144 | + | promoter-TSS (ENSMUSG00000029253) | -84 | Cenpc1 |
| Interval_30607 | 54,45130071 | -1,257058279 | 0,000212272 | 19 | 18619187 | 18620437 | + | intron (ENSMUSG00000024725, intron 1 of 10) | 11977 | Ostf1 |
| Interval_3816 | 51,29338043 | -1,278227131 | 0,000229145 | 1 | 173977544 | 173978232 | + | intron (ENSMUSG00000026535, intron 1 of 6) | 4856 | Ifi202b |
| Interval_1702 | 79,43648872 | 1,083395583 | 0,000231854 | 1 | 88213636 | 88214948 | + | intron (ENSMUSG00000089960, intron 1 of 4) | 2333 | Ugt1a1 |
| Interval_20339 | 188,585929 | 0,70617437 | 0,000249264 | 14 | 121937672 | 121939330 | + | intron (ENSMUSG00000041765, intron 1 of 3) | -7813 | Ubac2 |
| Interval_19401 | 34,94976441 | 1,658884581 | 0,00026309 | 14 | 69823411 | 69823912 | + | Intergenic | -16759 | Pebp4 |
| Interval_59140 | 36,41273826 | 1,56398437 | 0,000263996 | 9 | 61965345 | 61965995 | + | intron (ENSMUSG00000032278, intron 2 of 4) | -40549 | Kif23 |

# C

WT *vs* Δ3'RR in activated B cells

| PeakID | baseMean | log2FoldChange | pvalue | Chr | Start | End | Strand | Annotation | Distance to TSS | Gene Name |
|---|---|---|---|---|---|---|---|---|---|---|
| Interval_42073 | 233,3062839 | 1,864429221 | 1,03407E-06 | 4 | 118547560 | 118548718 | + | Intergenic | -4850 | Tmem125 |
| Interval_12753 | 33,5991101 | 2,150470647 | 5,34371E-05 | 12 | 31973306 | 31973947 | + | intron (ENSMUSG00000002997, intron 5 of 10) | -33219 | Hbp1 |
| Interval_1702 | 79,43648872 | 1,396831824 | 6,974E-05 | 1 | 88213636 | 88214948 | + | intron (ENSMUSG00000089960, intron 1 of 4) | 2333 | Ugt1a1 |

| | | | | | | | | | | |
|---|---|---|---|---|---|---|---|---|---|---|
| Interval_5472 | 132,8493816 | -1,208150001 | 0,000105676 | 10 | 44380319 | 44382062 | + | Intergenic | 21514 | Mir1929 |
| Interval_47232 | 37,14445921 | 1,960240443 | 0,00012695 | 5 | 137596735 | 137597133 | + | TTS (ENSMUSG00000037221) | 3757 | Mospd3 |
| Interval_10679 | 149,4549722 | -1,245289127 | 0,000140669 | 11 | 87442493 | 87444022 | + | promoter-TSS (ENSMUSG00000098943) | 20 | Rnu3b1 |
| Interval_59722 | 61,27660221 | -1,751427473 | 0,000154888 | 9 | 80057522 | 80058632 | + | Intergenic | -9740 | Senp6 |
| Interval_33995 | 187,3979067 | -1,0141174 | 0,000202174 | 2 | 90579301 | 90581549 | + | exon (ENSMUSG00000025314, exon 1 of 24) | 222 | Ptprj |

## D

WT *vs* c3'RR in activated B cells

| PeakID | baseMean | log2FoldChange | pvalue | Chr | Start | End | Strand | Annotation | Distance to TSS | Gene Name |
|---|---|---|---|---|---|---|---|---|---|---|
| Interval_1702 | 79,43648872 | 1,574730358 | 9,6632E-09 | 1 | 88213636 | 88214948 | + | intron (ENSMUSG00000089960, intron 1 of 4) | 2333 | Ugt1a1 |
| Interval_26244 | 38,10047802 | 2,209881998 | 1,96598E-08 | 17 | 30901814 | 30902460 | + | intron (ENSMUSG00000024027, intron 1 of 13) | 278 | Glp1r |
| Interval_42073 | 233,3062839 | 1,70172397 | 4,17253E-08 | 4 | 118547560 | 118548718 | + | Intergenic | -4850 | Tmem125 |
| Interval_21965 | 32,17992055 | 2,484747824 | 6,44651E-08 | 15 | 79900715 | 79901440 | + | TTS (ENSMUSG00000009585) | 735 | Apobec3 |
| Interval_35426 | 131,2968808 | -1,507463092 | 9,62783E-08 | 2 | 148763686 | 148764765 | + | Intergenic | 7272 | Cst11 |
| Interval_31829 | 42,87674911 | 2,155830932 | 1,16867E-07 | 19 | 61151514 | 61152486 | + | Intergenic | -11183 | Zfp950 |
| Interval_44094 | 45,62720152 | 2,071611603 | 5,10844E-07 | 5 | 14953946 | 14954607 | + | Intergenic | -15847 | Speer4e |
| Interval_44183 | 32,48779374 | 2,400333736 | 6,26839E-07 | 5 | 15674456 | 15675023 | + | Intergenic | -5971 | Speer4cos |
| Interval_22720 | 21,47947807 | 3,156788772 | 1,29761E-06 | 15 | 100443572 | 100443968 | + | Intergenic | -5914 | n-R5s43 |
| Interval_31323 | 44,29469979 | -1,768033742 | 3,19341E-06 | 19 | 44351580 | 44352475 | + | Intergenic | 18935 | Scd4 |
| Interval_19401 | 34,94976441 | 1,869601741 | 3,68295E-06 | 14 | 69823411 | 69823912 | + | Intergenic | -16759 | Pebp4 |
| Interval_3816 | 51,29338043 | -1,983993196 | 3,95923E-06 | 1 | 173977544 | 173978232 | + | intron (ENSMUSG00000026535, intron 1 of 6) | 4856 | Ifi202b |
| Interval_44187 | 132,6956757 | 1,818150838 | 5,11197E-06 | 5 | 15688990 | 15691064 | + | intron (ENSMUSG00000089871, intron 1 of 4) | 9317 | Speer4cos |
| Interval_44164 | 30,06497591 | 2,159268128 | 6,03011E-06 | 5 | 15612559 | 15613311 | + | intron (ENSMUSG00000094230, intron 2 of 2) | -7227 | Speer4d |
| Interval_44093 | 27,51772661 | 2,145276739 | 8,19656E-06 | 5 | 14952075 | 14952520 | + | Intergenic | -13868 | Speer4e |
| Interval_3614 | 11,9978808 | 6,648824762 | 1,01262E-05 | 1 | 171065212 | 171065430 | + | promoter-TSS (ENSMUSG00000059498) | -669 | Mir6546 |
| Interval_44188 | 22,32978783 | 2,395398899 | 1,16701E-05 | 5 | 15697839 | 15698169 | + | intron (ENSMUSG00000089871, intron 1 of 4) | 16267 | Speer4c |
| Interval_44186 | 113,2661305 | 1,853039056 | 1,38283E-05 | 5 | 15680019 | 15681848 | + | intron (ENSMUSG00000089871, intron 1 of 4) | 223 | Speer4cos |
| Interval_6230 | 124,1870108 | -1,071706257 | 2,06779E-05 | 10 | 75222061 | 75223178 | + | intron (ENSMUSG00000033444, intron 2 of 17) | -38348 | Specc1l |
| Interval_44184 | 40,29153996 | 1,730058639 | 2,76438E-05 | 5 | 15678270 | 15678776 | + | Intergenic | -2187 | Speer4cos |
| Interval_15282 | 105,7124407 | -0,992545515 | 2,88585E-05 | 13 | 21734066 | 21735952 | + | promoter-TSS (ENSMUSG00000067455) | -55 | H4c11 |
| Interval_22866 | 364,4687526 | -0,706606264 | 3,85548E-05 | 15 | 103239341 | 103241098 | + | promoter-TSS (ENSMUSG00000046434) | -275 | Hnrnpa1 |
| Interval_60870 | 711,7367054 | -2,705222962 | 4,33377E-05 | 9 | 123927808 | 123929581 | + | Intergenic | 39998 | Ccr1 |
| Interval_3611 | 55,66941374 | 1,519453318 | 4,50646E-05 | 1 | 171043984 | 171045076 | + | Intergenic | 14873 | Fcgr3 |
| Interval_47232 | 37,14445921 | 1,666946345 | 5,42141E-05 | 5 | 137596735 | 137597133 | + | TTS (ENSMUSG00000037221) | 3757 | Mospd3 |
| Interval_45797 | 131,6538692 | -0,897779869 | 5,51586E-05 | 5 | 96221157 | 96223265 | + | intron (ENSMUSG00000029486, intron 2 of 6) | 12067 | Mrpl1 |
| Interval_15347 | 82,63239193 | -1,068115974 | 8,69491E-05 | 13 | 23534826 | 23535728 | + | promoter-TSS (ENSMUSG00000099517) | -145 | H3c8 |
| Interval_30464 | 317,3291162 | -0,660957985 | 8,88432E-05 | 19 | 11964905 | 11966219 | + | promoter-TSS (ENSMUSG00000024687) | -379 | Osbp |

| | | | | | | | | | | |
|---|---|---|---|---|---|---|---|---|---|---|
| Interval_26224 | 26,54437711 | 1,982402618 | 9,46021E-05 | 17 | 30518826 | 30519215 | + | intron (ENSMUSG00000062202, intron 1 of 3) | 5883 | Btbd9 |
| Interval_34114 | 142,9101241 | -0,882283421 | 0,000107796 | 2 | 93427054 | 93428636 | + | intron (ENSMUSG00000027215, intron 6 of 6) | -5838 | Mir7001 |
| Interval_44302 | 13,78885203 | -2,562278039 | 0,000108172 | 5 | 21558544 | 21559035 | + | intron (ENSMUSG00000048520, intron 9 of 18) | 15230 | Lrrc17 |
| Interval_29172 | 243,8779389 | -0,779322032 | 0,000115394 | 18 | 60803129 | 60804071 | + | promoter-TSS (ENSMUSG00000024610) | -305 | Cd74 |
| Interval_21963 | 110,4203818 | 1,001117949 | 0,000142526 | 15 | 79891525 | 79893135 | + | promoter-TSS (ENSMUSG00000009585) | 2798 | AC113595.1 |
| Interval_21964 | 51,86372983 | 4,082461674 | 0,000153837 | 15 | 79894946 | 79896164 | + | exon (ENSMUSG00000009585, exon 2 of 8) | -4787 | Apobec3 |
| Interval_1712 | 25,20582771 | 1,918411199 | 0,00017025 | 1 | 88300647 | 88300934 | + | intron (ENSMUSG00000036251, intron 1 of 5) | -18123 | Trpm8 |
| Interval_14878 | 130,5275533 | -0,971499777 | 0,000170437 | 13 | 3525899 | 3526891 | + | Intergenic | -27644 | Gdi2 |
| Interval_15284 | 161,9871444 | -0,782875175 | 0,000172515 | 13 | 21753266 | 21754892 | + | promoter-TSS (ENSMUSG00000063021) | -44 | H2bc15 |
| Interval_26454 | 144,4452703 | -0,909899924 | 0,000189353 | 17 | 34257405 | 34258215 | + | exon (ENSMUSG00000073421, exon 1 of 3) | -5440 | H2-Ab1 |
| Interval_40863 | 103,4302046 | -1,078185572 | 0,000189918 | 4 | 53277689 | 53278635 | + | Intergenic | -12670 | AI427809 |
| Interval_41725 | 47,00441879 | -1,382843908 | 0,000190004 | 4 | 105263481 | 105264243 | + | Intergenic | 106515 | Plpp3 |
| Interval_50529 | 109,2901486 | -1,061151687 | 0,000206901 | 6 | 125572473 | 125573699 | + | intron (ENSMUSG00000001930, intron 6 of 14) | 20123 | Vwf |
| Interval_47180 | 64,90966257 | -1,140198511 | 0,000217993 | 5 | 136640538 | 136641571 | + | Intergenic | -56725 | Myl10 |
| Interval_44168 | 34,46301788 | 1,728339739 | 0,000228464 | 5 | 15632164 | 15632812 | + | intron (ENSMUSG00000094230, intron 2 of 2) | 12326 | Speer4d |
| Interval_40012 | 21,56651958 | -1,879694059 | 0,00023588 | 4 | 9640702 | 9641342 | + | intron (ENSMUSG00000028207, intron 1 of 4) | 2695 | Asph |
| Interval_3770 | 24,14270976 | 1,789781071 | 0,000245681 | 1 | 173490800 | 173491426 | + | promoter-TSS (ENSMUSG00000037849) | -17187 | Ifi206 |
| Interval_55548 | 45,58801231 | 1,545998902 | 0,000250773 | 8 | 33930207 | 33931049 | + | promoter-TSS (ENSMUSG00000031586) | -1236 | Rbpms |
| Interval_26245 | 11,73009879 | 2,714798357 | 0,00025438 | 17 | 30907467 | 30907703 | + | intron (ENSMUSG00000024027, intron 1 of 13) | 5726 | Glp1r |
| Interval_51254 | 323,8723748 | -0,69664298 | 0,000270884 | 7 | 3644569 | 3645690 | + | promoter-TSS (ENSMUSG00000035632) | -139 | Cnot3 |
| Interval_5472 | 132,8493816 | -0,813153144 | 0,000277818 | 10 | 44380319 | 44382062 | + | Intergenic | 21514 | Mir1929 |
| Interval_44088 | 30,47068406 | 1,776983366 | 0,000293044 | 5 | 14940262 | 14940887 | + | Intergenic | -2145 | Speer4e |
| Interval_1704 | 21,35051159 | 2,094514512 | 0,000297615 | 1 | 88222931 | 88223288 | + | Intergenic | -3877 | Mroh2a |
| Interval_1711 | 25,49836405 | 1,767036919 | 0,00030777 | 1 | 88292041 | 88292380 | + | intron (ENSMUSG00000036251, intron 1 of 5) | -21263 | Hjurp |
| Interval_6364 | 188,3314937 | -0,748373791 | 0,000311505 | 10 | 78086406 | 78087696 | + | Intergenic | 13712 | Icosl |
| Interval_6903 | 13,08088675 | 2,870172895 | 0,00032234 | 10 | 93198355 | 93198727 | + | intron (ENSMUSG00000020015, intron 1 of 2) | 35756 | Mir1931 |
| Interval_8250 | 13,00628344 | -3,595617214 | 0,00034183 | 11 | 6006357 | 6006741 | + | intron (ENSMUSG00000057897, intron 3 of 10) | -22411 | Camk2b |
| Interval_44167 | 23,28036799 | 1,865273345 | 0,000341887 | 5 | 15624001 | 15625093 | + | TTS (ENSMUSG00000070933) | 4385 | Speer4d |
| Interval_32299 | 216,176694 | -0,709439392 | 0,000346953 | 2 | 22773296 | 22775070 | + | promoter-TSS (ENSMUSG00000026786) | -207 | Apbb1ip |
| Interval_44181 | 18,0304766 | 2,268498764 | 0,000356485 | 5 | 15670992 | 15671420 | + | Intergenic | -9504 | Speer4cos |
| Interval_22557 | 63,60686106 | 1,253044281 | 0,000358304 | 15 | 97458206 | 97459223 | + | Intergenic | 97512 | Pced1b |
| Interval_3919 | 273,6291358 | -0,659543071 | 0,000390299 | 1 | 178186997 | 178188050 | + | promoter-TSS (ENSMUSG00000026502) | 106 | Desi2 |
| Interval_15370 | 295,3522521 | -0,628231787 | 0,000394584 | 13 | 23760742 | 23762873 | + | promoter-TSS (ENSMUSG00000060093) | -577 | H4c1 |
| Interval_15349 | 95,13161521 | -0,923886173 | 0,000408953 | 13 | 23550775 | 23552006 | + | exon (ENSMUSG00000069274, exon 1 of 1) | 258 | H4c6 |
| Interval_376 | 20,53100598 | 2,050888202 | 0,000449203 | 1 | 33883063 | 33883748 | + | promoter-TSS (ENSMUSG00000042182) | 75 | Bend6 |
| Interval_33989 | 71,50294514 | -1,153223244 | 0,000461867 | 2 | 90547072 | 90547800 | + | intron (ENSMUSG00000025314, intron 1 of 23) | 33211 | Ptprj |
| Interval_8579 | 88,94296747 | -0,972741454 | 0,000484739 | 11 | 22935942 | 22937170 | + | intron (ENSMUSG00000051355, intron 1 of 1) | 36303 | Commd1 |
| Interval_26229 | 208,4196308 | 1,309514607 | 0,000484765 | 17 | 30611699 | 30612947 | + | intron (ENSMUSG00000024026, intron 1 of 2) | 246 | Glo1 |

| | | | | | | | | | | |
|---|---|---|---|---|---|---|---|---|---|---|
| Interval_8283 | 30,33147907 | -1,831873649 | 0,000490909 | 11 | 6497445 | 6498063 | + | Intergenic | -21837 | Purb |
| Interval_37946 | 97,07367932 | -0,937381159 | 0,000491146 | 3 | 87401074 | 87402162 | + | intron (ENSMUSG00000059994, intron 10 of 10) | 25158 | Fcrl1 |
| Interval_15313 | 139,2061391 | -0,765686035 | 0,000506773 | 13 | 22035040 | 22036939 | + | promoter-TSS (ENSMUSG00000069302) | 119 | H2bc12 |
| Interval_57167 | 51,72714388 | -1,299461372 | 0,000515874 | 8 | 109750497 | 109751074 | + | Intergenic | -18672 | Atxn1l |
| Interval_51462 | 210,1594425 | -0,682504995 | 0,000530183 | 7 | 12977554 | 12978703 | + | promoter-TSS (ENSMUSG00000033961) | 235 | Zfp446 |
| Interval_59918 | 140,8688252 | -0,770390845 | 0,000541605 | 9 | 90059740 | 90061248 | + | exon (ENSMUSG00000032359, exon 2 of 12) | 5940 | Ctsh |
| Interval_23609 | 40,12752824 | -1,308519993 | 0,000567463 | 16 | 27399071 | 27400151 | + | intron (ENSMUSG00000038127, intron 1 of 8) | 10742 | Ccdc50 |

**Table S3 : Coordinates used for quantification of normalized coverage in R1, R2 and R3 regions**

| | chr12 | |
|---|---|---|
| **R1** | 113 216 000 | 113 225 000 |
| **R2** | 113 255 000 | 113 407 000 |
| **R3** | 113 407 000 | 113 430 000 |

**Table S4 : Normalized reads count of 3C-HTGTS performed with *Iµ/Sµ* bait in region of interest within the *IgH* locus in resting and stimulated B cells from *wt* and mutants mice**

**A**

| | chr12 | |
|---|---|---|
| **Sµ** | 113 385 000 | 113 430 000 |
| **Sγ3** | 113 355 000 | 113 370 000 |
| **Sγ2b** | 113 315 000 | 113 300 000 |
| **3'RR** | 113 255 000 | 113 385 000 |
| **eS** | 113 200 000 | 113 215 000 |
| **Whole IgH** | 113 200 000 | 133 430 000 |

**B**

| | **Resting B cells** | | | | | | | | | |
|---|---|---|---|---|---|---|---|---|---|---|
| | **WT#1** | **WT#2** | **WT#3** | **WT#4** | **WT#5** | **Δ3RR#1** | **Δ3RR#2** | **Δ3RR#3** | **c3'RR#1** | **c3'RR#2** |
| **Sµ** | 30 | 58 | 56 | 28 | 43 | 18 | 53 | 41 | 47 | 34 |
| **3'RR** | 109 | 252 | 195 | 76 | 100 | 33 | 132 | 143 | 134 | 107 |
| **eS** | 3538 | 3096 | 2802 | 4233 | 3867 | 3651 | 2445 | 3588 | 3015 | 4365 |
| **Whole IgH** | 1634 | 2325 | 2287 | 1136 | 1326 | 390 | 1100 | 992 | 1488 | 960 |

**C**

| | ***In vitro* stimulated B cells** | | | | | | | | | |
|---|---|---|---|---|---|---|---|---|---|---|
| | **WT#1** | **WT#2** | **WT#3** | **WT#4** | **Δ3RR#1** | **Δ3RR#2** | **Δ3RR#3** | **c3'RR#1** | **c3'RR#2** | **c3'RR#3** |
| **Sµ** | 82 | 68 | 50 | 40 | 15 | 11 | 5 | 82 | 47 | 61 |
| **Sγ3** | 1228 | 1247 | 1067 | 881 | 114 | 113 | 50 | 501 | 484 | 538 |
| **Sγ2b** | 169 | 178 | 129 | 94 | 10 | 25 | 15 | 74 | 135 | 102 |
| **3'RR** | 276 | 312 | 258 | 191 | 22 | 39 | 40 | 186 | 228 | 224 |
| **eS** | 2313 | 3043 | 3527 | 4214 | 4503 | 3459 | 4299 | 2643 | 4315 | 4076 |
| **Whole IgH** | 2892 | 2981 | 2430 | 1863 | 277 | 415 | 212 | 1552 | 1957 | 1900 |
